# Large-scale placenta DNA methylation mega-analysis reveals fetal sex-specific differentially methylated CpG sites and regions

**DOI:** 10.1101/2021.03.04.433985

**Authors:** Shan V. Andrews, Irene J. Yang, Karolin Froehlich, Tomiko Oskotsky, Marina Sirota

## Abstract

Although male-female differences in placental structure and function have been observed, little is understood about their molecular underpinnings. Here, we present a mega-analysis of 14 publicly available placenta DNA methylation (DNAm) microarray datasets to identify individual CpGs and regions associated with fetal sex. In the discovery dataset of placentas from full term pregnancies (N = 532 samples), 5,212 CpGs met genome-wide significance (p < 1E-8) and were enriched in pathways such as keratinization (FDR p-value = 7.37E-14), chemokine activity (FDR p-value = 1.56E-2), and eosinophil migration (FDR p-value = 1.83E-2). Nine differentially methylated regions were identified (fwerArea < 0.1) including a region in the promoter of *ZNF300* that showed consistent differential DNAm in samples from earlier timepoints in pregnancy and appeared to be driven predominately by effects in the trophoblast cell type. We describe the largest study of fetal sex differences in placenta DNAm performed to date, revealing genes and pathways characterizing sex-specific placenta function and health outcomes later in life.

## Introduction

The placenta is a key organ during pregnancy, performing important functions such as providing nutrients, transferring respiratory gases, and secreting hormones for adequate fetal growth and development^1^. During pregnancy, the placenta grows and changes in composition and function, developing within the mother, but is primarily regulated by the fetal genome^1,2^. Epidemiologic evidence suggests that the periconception and *in utero* periods are the most vulnerable to environmental factors influencing the susceptibility for several diseases later in life^3,4^. The placenta, as the key mediator of the gestational environment, therefore has a significant role in the fetal programming of health outcomes into adulthood^5^. Further, observed differences by sex with respect to prevalence or severity in these health outcomes, as has been observed for autism spectrum disorder (ASD)^6^, schizophrenia^7^, and autoimmune diseases^8^, among many other examples, may be influenced by sex-specific gestational environments that are mediated by sex-specific placenta structure and function.

Some male-female differences during *in utero* fetal development and childbirth are already known. For example, male fetuses are characterized by higher birth weight, placenta weight, and birth weight to placenta weight ratio^9,10^. Moreover, males have a higher risk of suffering from peri- and postnatal complications^11–16^. Previous studies have demonstrated that male and female fetuses utilize different mechanisms to cope with adverse intrauterine environments^17–20^. For instance, in the presence of chronic maternal asthma growth of female fetuses is reduced, resulting in neonates of smaller size and lower birth weight compared to neonates from healthy pregnancies^18,21^. Any further complication during pregnancy, such as an acute maternal asthma exacerbation, did not decrease their survival rate, indicating a vital adaptation mechanism of female fetuses^21,22^. In contrast, growth of male fetuses appears not to be reduced in the presence of maternal chronic asthma; however, male fetuses experienced a higher rate of adverse outcomes among women who had a severe asthma exacerbation^18,20^.

A growing body of literature has sought to understand the placental molecular mechanisms at play in disease outcomes or complicated pregnancies. Placenta gene expression studies have examined differences in preeclampsia^23,24^, gestational diabetes mellitus^25^, and intrauterine growth restriction (IUGR)^26^. For DNA methylation (DNAm), examples of studies include those examining preterm birth^27^, ASD^28^, and also preeclampsia^29^ and IUGR^30^. There have been some examples in these studies of analyses that explicitly seek to quantify sex-specific mechanisms. For example, gene expression differences, including for various cytokines, have been shown to characterize fetal sex-specific responses to maternal asthma exposure^31–33^. However, these sex-focused analyses are rare, and there have been few studies that have examined sex itself as an “outcome” in analyses, even in normal or non-pathological placentas. Finally, the majority of transcriptomic and DNAm studies of the placenta have not thoroughly accounted for or investigated the role of cell type heterogeneity in phenotypic studies^34^, though a recent study by Yuan et al.^35^ has helped to address this gap by generating DNAm profiles in the four major placenta cell types. Specifically, this study profiled endothelial cells, placental macrophages known as Hofbauer cells, mesenchymal stromal cells and trophoblasts, composed of both syncytiotrophoblasts that interact with maternal blood and cytotrophoblasts that invade the uterine spiral arteries^36^. Overall, the mechanisms by which placental molecular factors act in a sex-specific manner to determine fetal phenotypes, and how these mechanisms act through specific cell types or change with gestational age, remain as critical gaps in our understanding of developmental health. While placental gene expression studies investigating sex-specific differences have been undertaken^37–40^, previous studies of placenta DNAm related to fetal sex^41,42^ have suffered from small sample size.

Here, we present a mega-analysis of publicly available placenta DNAm samples from uncomplicated pregnancies in order to maximize sample size for identification of sex-specific differences. We first sought to characterize sex-specific patterns in DNAm in full term placentas, aiming to identify differentially methylated CpG sites as well as differentially methylated regions (DMRs). We then performed similar analyses using placenta samples from earlier points in pregnancy (1st, 2nd, and 3rd trimesters individually) and evaluated which findings from full term placentas persist across the gestational period. Finally, we examined our results in light of recently published cell type specific placenta DNAm patterns, to understand the degree to which cell type heterogeneity drove our findings and if these patterns could shed light on the mechanism of our identified sites and regions of interest.

## Results

### Sex-specific methylation patterns of non-pathological human placenta tissue

We analyzed sex differences in human placenta DNAm patterns incorporating 783 samples from 14 Illumina 450K array GEO data sets (**Figure 1A, Supplementary Table 1**). Samples from control and/or normal pregnancies were binned into groups according to gestational period. The full term dataset was treated as the discovery dataset, owing to its large sample size. The smaller, less-statistically powered individual trimester datasets were treated as replication cohorts examining the extent to which differentially methylated sites and regions identified at full term persisted throughout the gestational period. In the full term dataset and specific trimester datasets, individual CpG sites and regions were evaluated for their association with fetal sex, adjusting for dataset-specific surrogate variables (see Methods). We verified that the surrogate variables captured batch effects resulting from the many individual studies that contributed to the datasets (**Figure 1B, Supplementary Figures 1-3**).

**Table 1:** Significant (FDR p-value < 0.05) Gene Ontology pathways enriched in differentially methylated sites from the full term placenta analysis.

| GO ID | ONTOLOGY | TERM | FDR p-value |
| --- | --- | --- | --- |
| *GO:0018149* | BP | peptide cross-linking | 7.37E-14 |
| *GO:0031424* | BP | keratinization | 7.37E-14 |
| *GO:0030216* | BP | keratinocyte differentiation | 4.07E-13 |
| *GO:0001533* | CC | cornified envelope | 1.93E-12 |
| *GO:0070268* | BP | cornification | 2.95E-12 |
| *GO:0009913* | BP | epidermal cell differentiation | 7.78E-12 |
| *GO:0043588* | BP | skin development | 3.64E-08 |
| *GO:0008544* | BP | epidermis development | 1.20E-07 |
| *GO:0045095* | CC | keratin filament | 1.67E-06 |
| *GO:0030855* | BP | epithelial cell differentiation | 2.41E-04 |
| *GO:0060429* | BP | epithelium development | 8.68E-04 |
| *GO:0099512* | CC | supramolecular fiber | 3.50E-03 |
| *GO:0099081* | CC | supramolecular polymer | 4.55E-03 |
| *GO:0099080* | CC | supramolecular complex | 4.55E-03 |
| *GO:0048020* | MF | CCR chemokine receptor binding | 1.12E-02 |
| *GO:0048245* | BP | eosinophil chemotaxis | 1.39E-02 |
| *GO:0008009* | MF | chemokine activity | 1.56E-02 |
| *GO:0072677* | BP | eosinophil migration | 1.83E-02 |
| *GO:0048513* | BP | animal organ development | 3.51E-02 |

**Figure 1.**
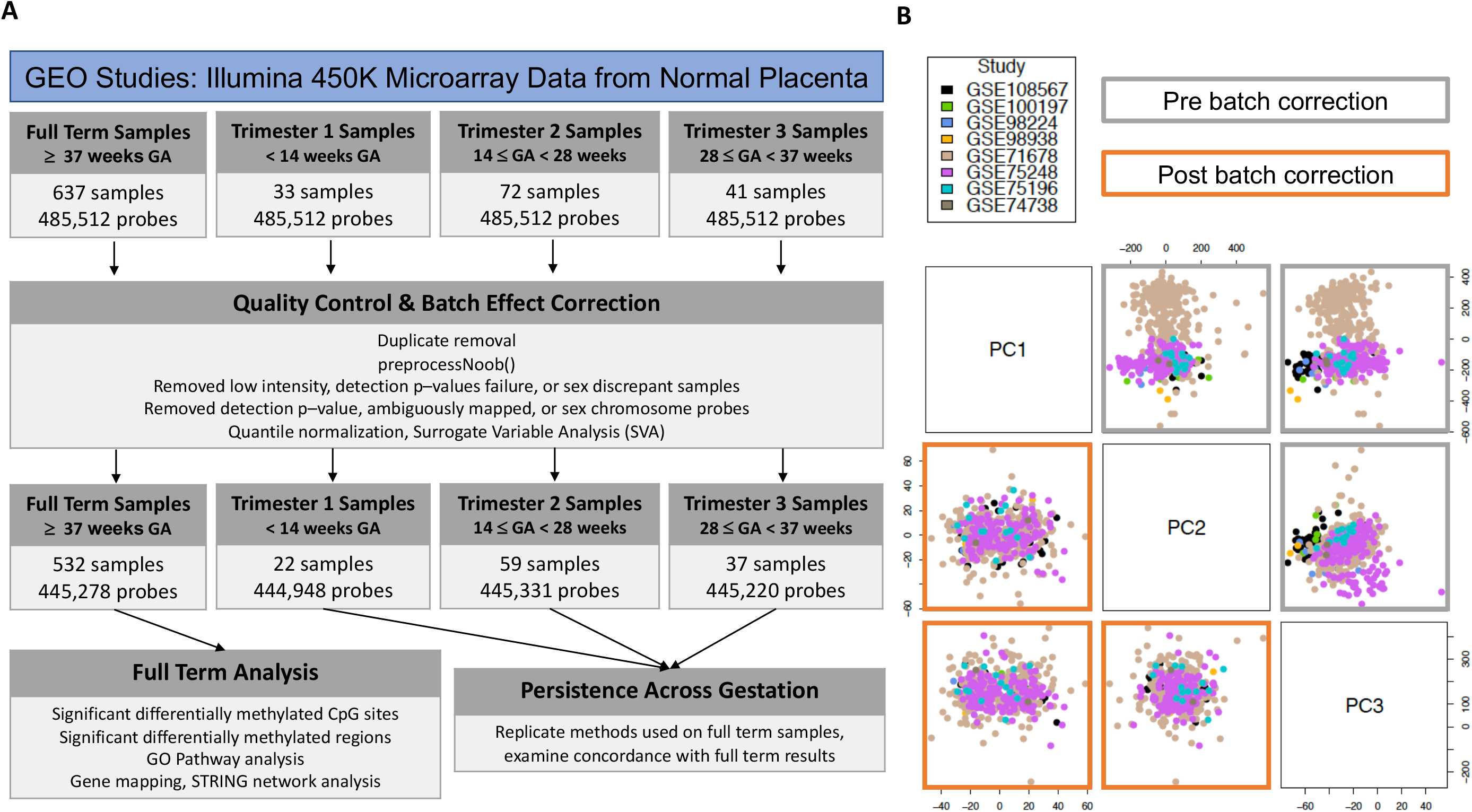
Methods overview and batch effect correction. **a** Illumina 450K datasets of fetal placenta samples were downloaded from GEO and restricted to those from normal/typical pregnancies. Samples were classified into 4 groups according to annotated gestational age, and each dataset was processed through identical subject and probe level QC steps. Single site and region-based analyses were performed in full term samples first, and then also performed in individual trimester datasets to examine replication of full term findings. See Methods for additional details. **b** Principal component plot depicting DNAm M-values (top triangle) and residuals from model of M-values regressed onto estimated surrogate variables (bottom triangle) in full term dataset. Points colored by GEO ID.

### Differentially methylated CpG sites by fetal sex in full term placenta samples

Single site analysis at 445,278 autosomal in 532 full term placenta samples (after QC; see Methods) revealed that a total of 5,212 CpG sites were significantly associated with fetal sex at a genome wide significance threshold of 1E-8 (**Figure 2A, Supplementary Data 1**). Of these, cg01382982, harbored in the CpG island promoter of *ZNF175*, exhibited the largest degree of hypermethylation in males (**Figure 2B**; mean difference = 22%, p-value = 9.12E-37). In females, the CpG site achieving genome-wide significance with the highest degree of hypermethylation was cg22905511 (**Figure 2C**; mean difference = -11%, p-value = 2.85E-13), which is located in a CpG island promoter of *C5orf63*. The majority (3,793; 73%) of genome-wide significant sites were hypermethylated in male samples; these male-hypermethylated sites tended to exhibit larger effect sizes than those hypermethylated in females (**Figure 2D**). Finally, genome-wide significant CpG sites were enriched in GO pathways involved in keratinization, cell differentiation as well as immune cell migration and chemotaxis (**Table 1, Supplementary Data 2**). There was very minimal overlap between the identified differentially methylated sites and published 450K-based genome-wide screens of sex-associated DNAm, across a variety of tissue types (**Table 2**). Though these studies varied in their methodological approach and probe filtering strategies, this result supports the conclusion that the majority of differentially methylated sites identified herein are placenta-specific.

**Table 2:** Overlap of significant findings from previous studies of differential DNAm by sex with full term placenta results. Results from previous studies restricted to autosomes. ^a^Percentage indicates proportion of significant sites from specified study included in placenta full term, post QC dataset. ^b^Percentage indicates proportion of significant sites from specified study found in significant findings from full term analysis (p < 1E-8, n = 5,212).

| Study | Year | Tissue | # Significant autosomal CpG sites differential by sex (Count included in full term, post QC analysis) | # Significant sites included in post QC, full term placenta dataset (%)<sup>a</sup> | Overlap with placenta full term results (%)<sup>b</sup> |
| --- | --- | --- | --- | --- | --- |
| Price et al. | 2013 | Peripheral blood, adult | 45 | 40 (88.89%) | 13 (0.25%) |
| Hall et al. | 2014 | Pancreatic islets | 470 | 339 (72.13%) | 55 (1.06%) |
| Xu et al. | 2014 | Prefrontal cortex | 13,560 | 13,554 (99.96%) | 331 (6.35%) |
| Inoshita et al. | 2015 | Peripheral leukocytes | 292 | 292 (100%) | 103 (1.98%) |
| Singmann et al. | 2015 | Peripheral blood | 11,010 | 11,008 (99.98%) | 530 (10.2%) |
| Spiers et al. | 2015 | Fetal cortex | 521 | 521 (100%) | 103 (1.98%) |
| Yousefi et al. | 2015 | Cord blood | 3,031 | 3,031 (100%) | 318 (6.10%) |
| Martin et al. | 2017 | Placenta | 15 | 1 (6.7%) | 1 (0.02%) |
| Suderman et al. | 2017 | Cord blood | 11,965 | 11,961 (99.97%) | 274 (5.26%) |
|  |  | Peripheral blood, 7 years | 13,672 | 13,669 (99.98%) | 310 (5.95%) |
|  |  | Peripheral blood, 15-17 years | 12,215 | 12,214 (99.99%) | 291 (5.58%) |
| Xia et al. | 2019 | Prefrontal cortex, Bulk Brain | 15,417 | 15,048 (97.61%) | 439 (8.42%) |

**Figure 2.**
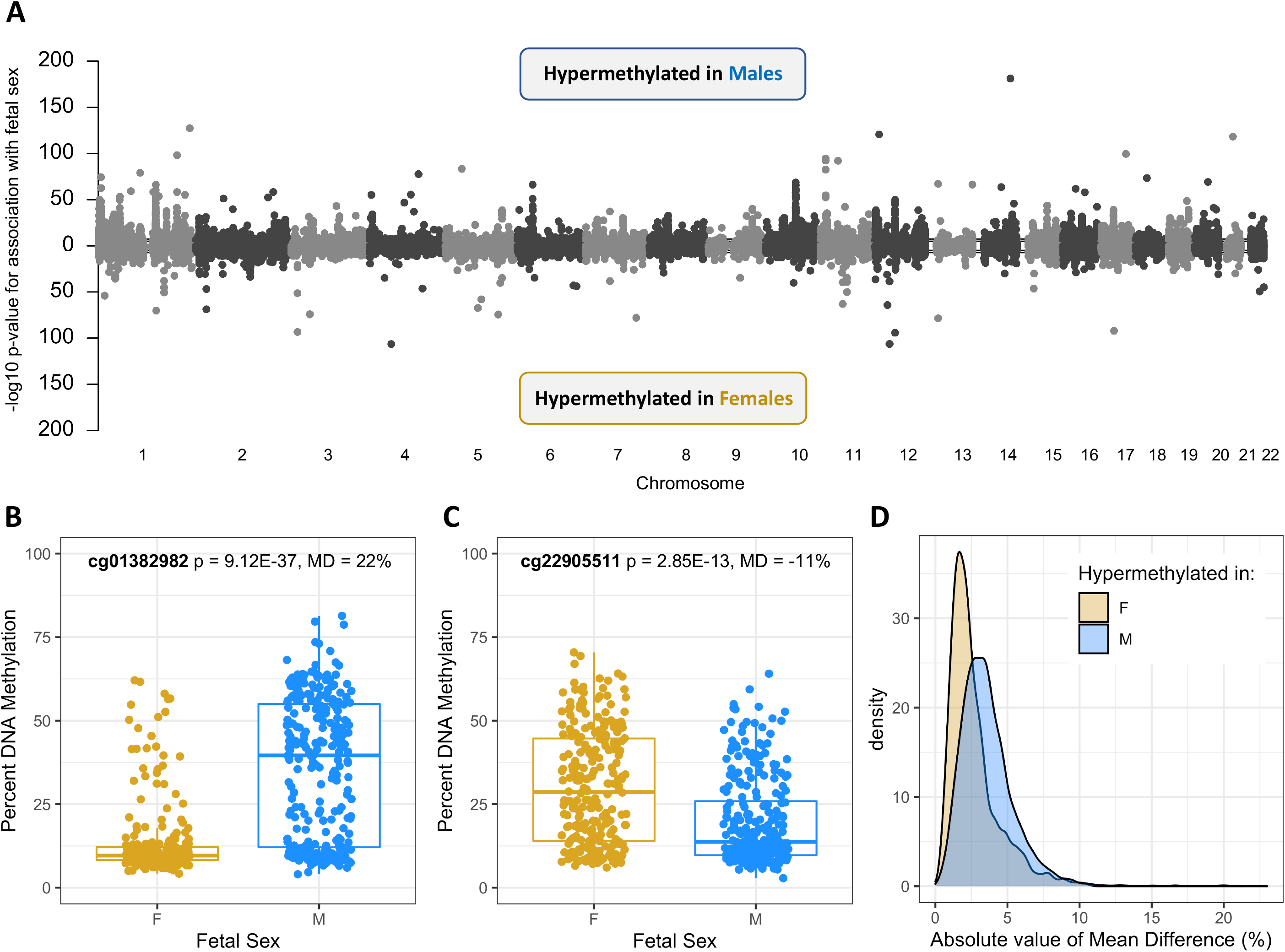
Single site plots. **a** The manhattan plot provides a global view of the CpG sites from the full term samples. The x-axis represents genomic position, and the y-axis represents the -log10 of the association’s p-value. Each point is a CpG site. **b** Boxplots of percent methylation in males (blue) and females (gold) at cg01382982, the genome-wide significant CpG site at which males were most hypermethylated relative to females (Mean Difference, MD = 22%). **c** Boxplots of percent methylation in males (blue) and females (gold) at cg22905511, the genome-wide significant CpG site at which females were most hypermethylated relative to males (Mean Difference, MD = -11%). **d** Distribution of absolute value of mean differences in sites hypermethylated in males (blue, n = 3,793) and sites hypermethylated in females (gold, n = 1,419).

Of the 5,212 differentially methylated sites, 1,285 (24.7%) were identified as distinguishing full term placenta cell types in a recent paper by Yuan et al. examining cell type specific DNAm patterns in term and early term placentas^35^. This subset of cell type distinguishing probes did not appear to differ markedly from those that did not distinguish cell type in the distribution of effect sizes from the fetal sex comparison (**Supplementary Figure 4**). However, GO pathway analysis stratified by probe group type revealed that while keratinization and cell differentiation pathways were shared in the two probe groups, the cell type distinguishing probes were uniquely enriched in chemokine and immune cell migration pathways (**Supplementary Data 3 and 4**).

### Persistence of differentially methylated CpG sites across the gestational period

We additionally identified publicly available DNAm samples from preterm placentas, and conducted trimester-specific QC and association analysis using identical methods as used in the full term samples (**Supplementary Figures 1-3**; see Methods for classification of placentas into these trimesters). The goal of these analyses of early gestation samples was to understand the extent to which differentially methylated sites identified in full term placenta samples persisted across gestation. Gestational age prediction verified that the full term and trimester-specific datasets (N = 22, 59, and 37 for Trimesters 1-3, respectively) captured their intended gestational periods (**Figure 3A**). Intuitively, greater correlation was observed in effect sizes as trimesters approached full term; for example the correlation between first trimester and full term effect sizes was 34% while the correlation between third trimester and full term effect sizes was 80% (**Figure 3B**). Of the 5,209 genome-wide significant CpG sites from the full term analysis (3 CpG sites did not pass QC in ≥ 1 trimester-specific dataset), 194 CpG sites replicated their association with fetal sex in each trimester (**Figure 3C**; replication defined as p-value ≤ 0.05 and consistent direction of effect as seen in full term samples). An example of one of these cross-gestation persistent CpG sites is cg17612569, which maintains a hypermethylated state in male placenta samples of at least 13% from the first trimester through full term (**Table 3, Figure 3D**). A protein-protein interaction network derived from the 194 persistent sites revealed a greater cluster (red) mainly associated to neurogenesis, axonogenesis, gliogenesis and developmental processes and two smaller clusters (green, light green) mainly associated to keratinization and peptide cross-linking. All other networks included less than four gene symbols (**Figure 3E**). Full genome-wide association statistics for the analyses from each trimester have been made available (**Supplementary Data 5-7**; **Supplementary Figure 5**).

**Table 3:** Top 10 CpG sites from full term analysis (ranked by absolute value of effect size in full term analysis) displaying persistence across the gestational period. Mean Difference (MD) values calculated as mean in male samples - mean in female samples.

| **CpG Site** | **MD, Full Term** | **P-value, Full Term** | **MD, 3rd Trimester** | **P-value, 3rd Trimester** | **MD, 2nd Trimester** | **P-value, 2nd Trimester** | **MD, 1st Trimester** | **P-value, 1st Trimester** |
| --- | --- | --- | --- | --- | --- | --- | --- | --- |
| cg02375258 | 20.4 | 1.46E-31 | 9.1 | 0.0387 | 14 | 0.00378 | 24.1 | 0.000743 |
| cg18237551 | 20 | 8.15E-35 | 19.8 | 0.000453 | 22.4 | 1.02E-05 | 18.4 | 0.00035 |
| cg08580836 | 19.9 | 9.05E-39 | 18.6 | 0.000253 | 26.3 | 2.83E-06 | 28.2 | 2.03E-05 |
| cg21228005 | 19.6 | 3.4E-39 | 19.6 | 0.00185 | 24.5 | 1.85E-06 | 27.6 | 2.1E-06 |
| cg02343823 | 18.2 | 3.58E-36 | 14.4 | 0.000688 | 18.6 | 1.2E-05 | 17.5 | 3.02E-05 |
| cg04675542 | 16.5 | 6.38E-30 | 13.3 | 0.00038 | 20.4 | 1.11E-05 | 16.4 | 5.69E-05 |
| cg11291313 | 13.4 | 1.01E-33 | 12 | 0.00164 | 18.2 | 5.29E-05 | 20.4 | 6.01E-05 |
| cg19014419 | 13 | 2.88E-34 | 13.4 | 0.000429 | 17.8 | 4.48E-05 | 18.6 | 0.000183 |
| cg17612569 | 12.6 | 5.93E-119 | 20 | 1.5E-13 | 20.7 | 3.5E-20 | 13.7 | 1.72E-07 |
| cg02325951 | 12.1 | 5.81E-182 | 13.4 | 3.14E-16 | 11.6 | 1.5E-12 | 11.8 | 1.17E-05 |

**Figure 3.**
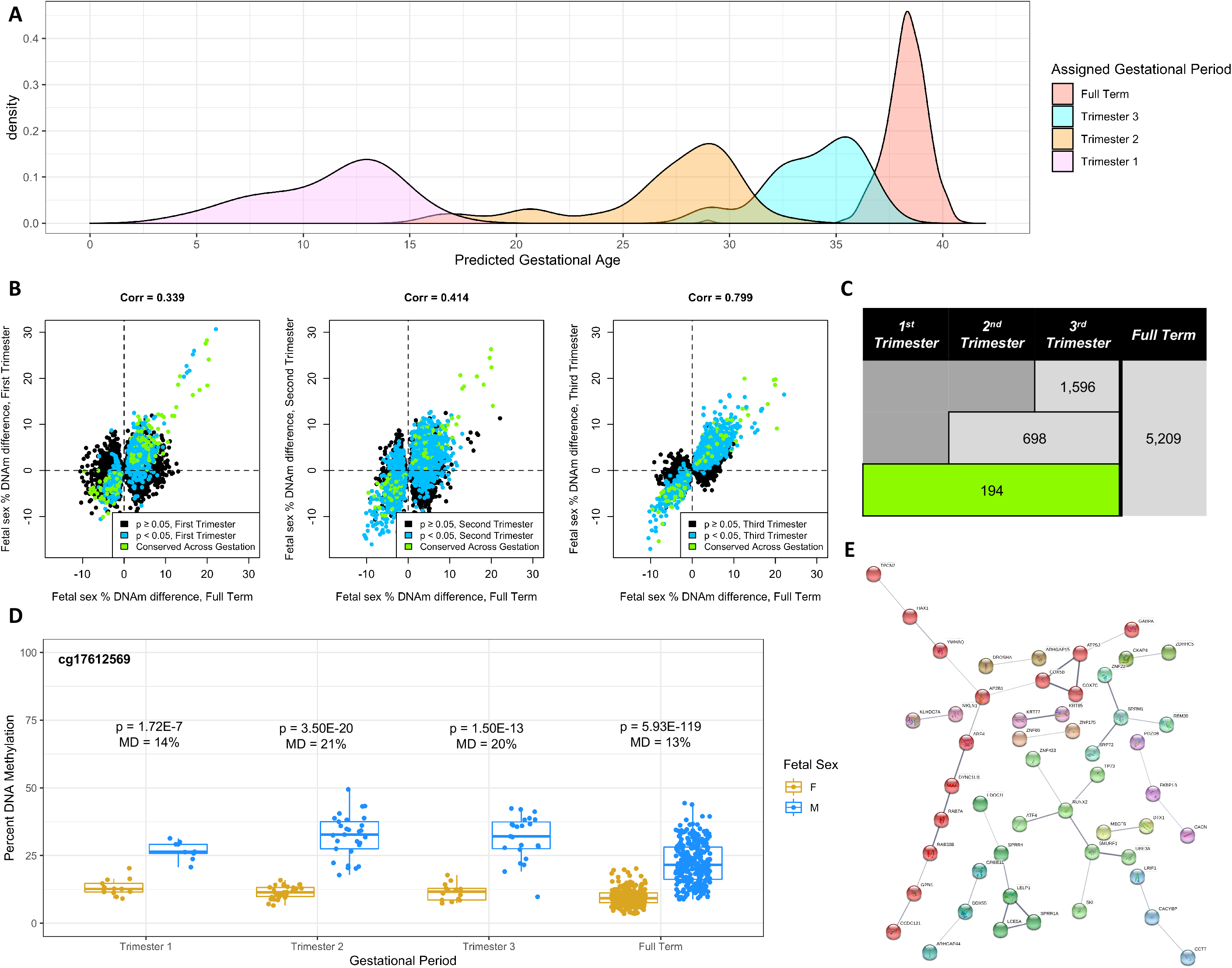
Persistence across the gestational period. **a** Distribution of predicted gestational age (via *predictAge()* function from the *planet* R package) in full term and individual trimester datasets **b** The persistence correlation analysis shows plots of DNA methylation differences for the three trimesters. The green colored points are the CpG sites that were conserved across gestation. The blue colored points represent the CpG sites that were considered significant for each trimester. The correlation values between the trimester and the full term analysis increase as the full term is approached. **c** Persistence of hits discovered from the full term placentas throughout gestation. If the significant site from the full term had a p value less than 0.05 and had the same direction of association in the preterm analysis, then it was considered “persistent.” **d** Comparison of female and male fetuses’ methylation values at different points during gestation. cg17612569 was consistently significant at all time points. **e** STRING network analysis based on genes mapped from persistent single sites shows protein-protein interactions. Each node is a protein. The higher confidence scores for the interactions are represented by the darker lines.

### Region-based differential DNAm analyses and persistence across gestation

We also sought to identify DMRs in full term samples and examine the extent to which they persisted across gestation. Nine DMRs were identified as significant beyond a discovery-based threshold (fwerArea value) of 0.1 (**Supplementary Figure 6, Supplementary Data 8**). The top-ranked DMR was located in a CpG island promoter of *ZNF175* (fwerArea = 0); males exhibited 17% higher average DNAm relative to females in this region (**Figure 4A**). Trimester-specific region-based analyses (**Supplementary Data 9-11**) were not statistically powered for genome-wide detection. However, these analyses did reveal that the identified *ZNF175* promoter region exhibited a similar degree of hypermethylation in males across the gestational period (**Figure 4B-D**). In fact, all of the nine full term DMRs exhibited evidence for persistence across gestation via consistent direction of effect across all trimesters (**Table 4, Supplementary Figures 7-14**). However, in most cases there was a significantly decreased magnitude of effect size in the individual trimesters. The most robust evidence for persistence, beyond the *ZNF175* DMR, was observed for male hypermethylation at *ZNF300* promoter (**Supplementary Figure 7**) and for female hypermethylation at the *C5orf63* promoter (**Supplementary Figure 8**). In the latter case, the DMR did not emerge until the 2nd trimester, but persisted thereafter at a similar magnitude.

**Figure 4.**
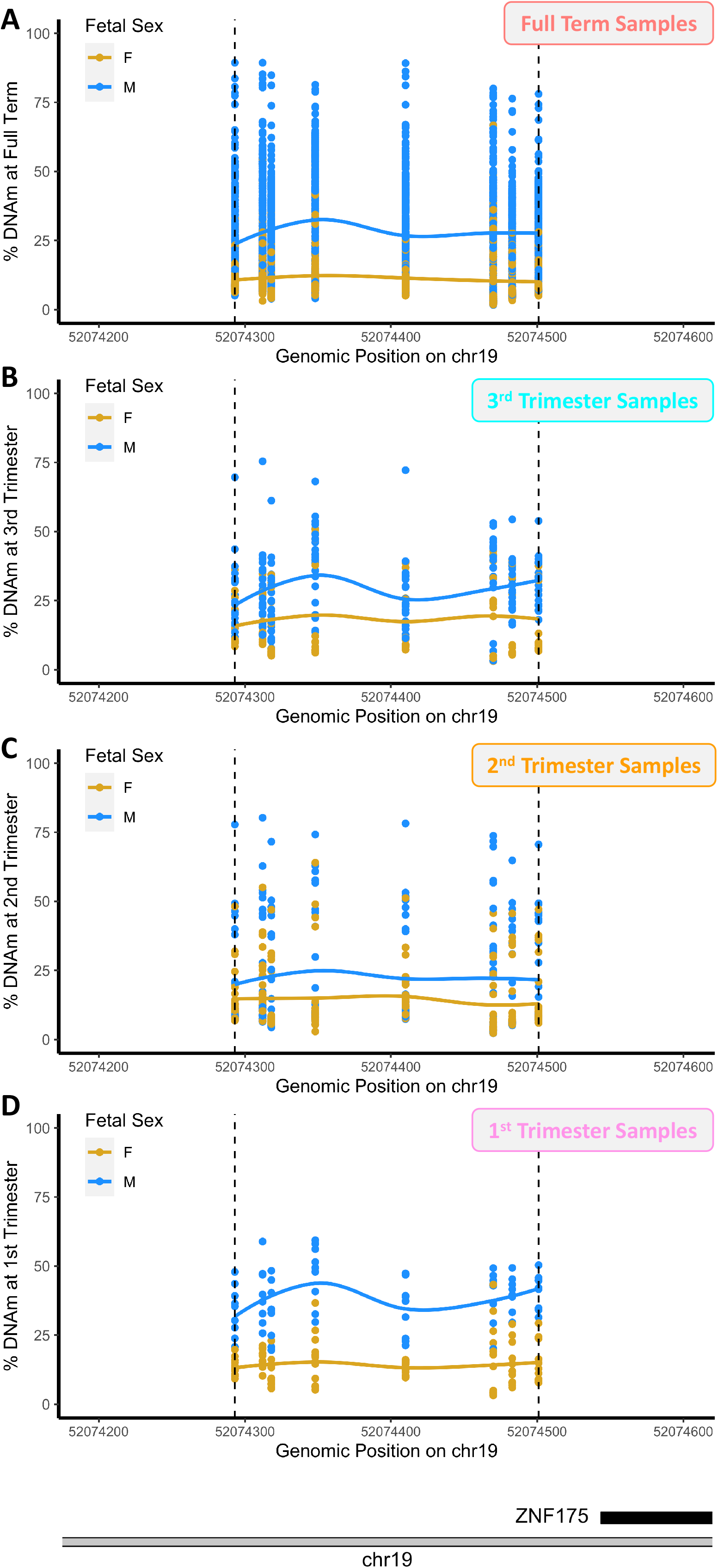
DMR at the *ZNF175* promoter across the gestational period. Top-ranked DMR from full term analysis at *ZNF175* promoter. DMRs are plotted as percent methylation as a function of genomic position. Dots indicate samples and solid lines indicate smooth lines through male sample values (blue) and female sample values (gold). Actual identified DMR region indicated via dashed lines. **a** Full Term Samples. **b** 3rd Trimester Samples **c** 2nd Trimester Samples **d** 1stTrimester Samples.

**Table 4:** Persistence of genome-wide significant differentially methylated regions from full term analysis across the gestational period. Statistics generated from *bumphunter()*^62^ function in minfi package^55^. Mean Difference (MD) values calculated as mean in male samples - mean in female samples. Final column indicates if DMR contained cell type distinguishing probe as identified by Yuan et al. in full term samples.

| chr | start | end | p.value | fwer | p.value Area | fwer Area | MD, Full Term | Nearest Gene (location) | MD, Trimester 1 | MD, Trimester 2 | MD, Trimester 3 | Contains cell type discriminating probes? |
| --- | --- | --- | --- | --- | --- | --- | --- | --- | --- | --- | --- | --- |
| chr19 | 52074293 | 52074501 | 0 | 0 | 0 | 0 | 0.17 | *ZNF175* (promoter) | 0.24 | 0.0831 | 0.11 | No |
| chr5 | 150284416 | 150284796 | 0 | 0 | 0 | 0 | 0.172 | *ZNF300* (promoter) | 0.21 | 0.212 | 0.159 | Yes |
| chr5 | 126408756 | 126409553 | 0 | 0 | 0 | 0 | -0.0737 | *C5orf63* (promoter) | -0.00439 | -0.101 | -0.101 | Yes |
| chr11 | 2906285 | 2907127 | 0 | 0 | 1.83E-06 | 0.007 | 0.0243 | *CDKN1C* (gene body) | 0.0325 | 0.0159 | 0.0295 | No |
| chr7 | 86974674 | 86975244 | 2.61E-07 | 0.001 | 3.66E-06 | 0.014 | -0.0522 | *CROT* (promoter) | -0.0514 | -0.0743 | -0.0761 | Yes |
| chr7 | 27169674 | 27171051 | 5.23E-07 | 0.002 | 1.05E-05 | 0.039 | -0.0372 | *HOXA4* (promoter) | -0.0106 | -0.0035 | 0.00492 | Yes |
| chr4 | 90758120 | 90759203 | 1.05E-06 | 0.004 | 1.73E-05 | 0.065 | -0.0393 | *SNCA* (promoter) | -0.0216 | -0.0138 | -0.00186 | No |
| chr6 | 28641622 | 28642394 | 3.4E-06 | 0.013 | 1.83E-05 | 0.069 | 0.0329 | *ZBED9* (58 Kb downstream) | 0.0419 | 0.0692 | 0.0452 | Yes |
| chr9 | 98637332 | 98638444 | 2.61E-07 | 0.001 | 2.25E-05 | 0.085 | -0.0561 | *ERCC6L2* (promoter) | -0.0427 | -0.0143 | -0.0321 | No |

Of the nine DMRs, five contained probes that were identified as distinguishing cell type by Yuan et al. (**Table 4**). Plotting the placenta cell type specific DNAm patterns from this study in the regions defining these DMRs revealed in several instances that particular cell types drive the fetal sex DNAm patterns we observed. For example, male hypermethylation in the *ZNF300* promoter is primarily driven by sex differences in trophoblast cell types in this region (**Figure 5A-B**). Just as we observed this fetal sex differential DNAm to persist throughout gestation (**Figure 5C, Table 4, Supplementary Figure 7**), the trophoblast signatures remain when examining the 1st trimester samples from Yuan et al. (**Figure 5D**). For the remaining four DMRs containing cell type distinguishing probes, cell type-related inferences are less clear (**Supplementary Figures 15-18**), likely owing to the smaller magnitude of DNAm difference in the DMRs. Nonetheless, it can be observed that sex differences in trophoblast DNAm contribute to the DMRs observed at the *C5orf63* (**Supplementary Figure 15**) and *CROT* (**Supplementary Figure 16**) promoters, with endothelial cells playing a role in the latter case as well, but only in the early gestation time point.

**Figure 5.**
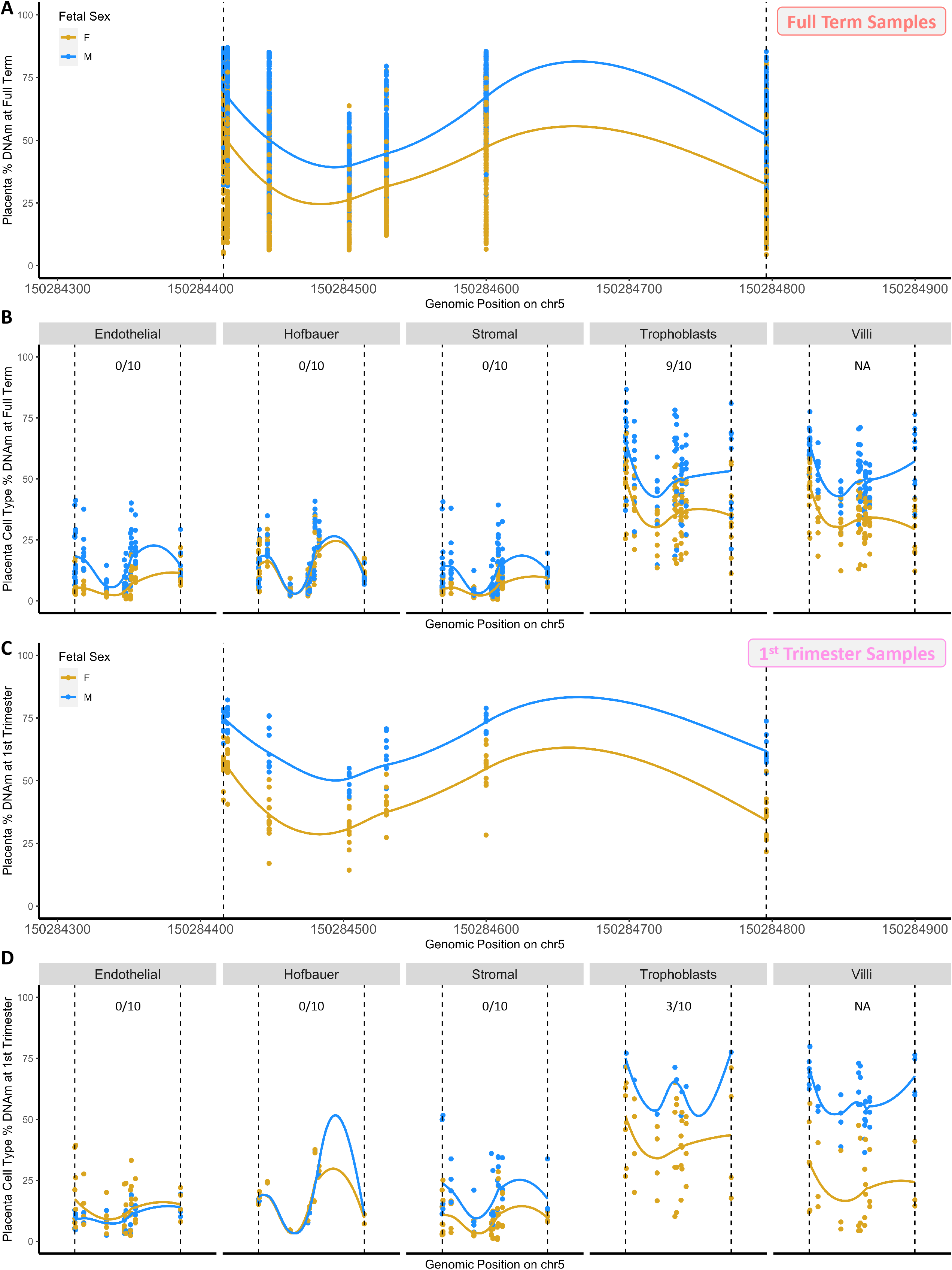
Cell type heterogeneity at the *ZNF300* promoter. DMRs are plotted as percent methylation as a function of genomic position. Dots indicate samples and solid lines indicate smooth lines through male sample values (blue) and female sample values (gold). Actual identified DMR region indicated via dashed lines. **a** DMR identified at *ZNF300* promoter on chromosome 5 in full term analysis, plotted using full term samples from present study. **b - d** The same region, but plotted using samples from different datasets. **b** Term samples and individual placenta cell types from Yuan et al. **c** 1st Trimester samples from present study. **d** 1st trimester samples and individual placenta cell types from Yuan et al. In **b** + **d**, fractions indicate the proportion of probes in the region that were annotated by Yuan et al. as distinguishing that cell type in full term and 1st trimester samples, respectively. Villi samples were not evaluated in this manner by Yuan et al. as this is the unsorted bulk tissue.

## Discussion

Here we present the largest study to date examining placental DNAm differences by fetal sex. Both single site and region-based analyses revealed large differences according to fetal sex in the placenta methylome. Single site analysis of full term placenta samples demonstrated that a total of 5,212 CpG sites were significantly associated with fetal sex at a genome-wide significance threshold of 1E-8. Pathway analysis results showed that some of the most significantly enriched biological processes were keratinization, cell differentiation, and eosinophil chemotaxis. Replication studies using samples at earlier points in gestation demonstrated evidence that 194 of these CpG sites are differentially methylated by sex throughout pregnancy. Nine DMRs met a discovery-based threshold of significance, and the top-ranked DMRs at the promoters of *ZNF175* and *ZNF300*, respectively, displayed robust evidence for persistence across the gestational period. In the latter case, cell type specific placenta DNAm data revealed that trophoblast cell types drive the fetal sex association observed in the region.

Our findings of many differentially methylated sites and regions according to fetal sex will contribute to the understanding of the molecular mechanisms driving sex-specific placenta structure and functions. For example, several immune-related pathways were enriched in differentially methylated CpG sites from the full term analysis, which is consistent with several previous investigations of sex-related gene expression differences in placenta^37,39,40^. A recent dataset described by Yuan et al^35^, which profiled DNAm in sorted placenta cell types, allowed for further characterization of these immune pathway findings. While the Yuan et al. study found no significant differences in estimated cell type proportions between male and female placentas, it is still possible that individual probes exhibit both sex-specific and cell type specific DNAm patterns. Indeed, we found that a significant proportion (1,285, 24.7%) of our 5,212 fetal sex differentially methylated sites in full term placentas were defined by Yuan et al. to distinguish placenta cell types at full term as well. Stratified pathway analyses considering probes in this group of 5,212 that did and did not distinguish placenta cell types separately revealed that the enrichment of chemokine and immune cell migration pathways seen in the overall group of probes was observed uniquely in the cell-type-distinguishing subset. This result argues that sex-specific immune responses in the placenta are mediated through cell type proportion changes.

Keratinization and cell differentiation pathways were also strongly enriched in the differentially methylated sites at full term, and were enriched in both the cell type distinguishing and non-cell-type-distinguishing probe subsets separately. Similar pathways were found in enrichment studies from a previous investigation of gene expression differences by fetal sex^38^. These pathways are also remarkably consistent with those previously found to be enriched in a gene set consisting of “core” placenta genes that are conserved across evolution and described as “central for making a placental mammal”^43^, indicating that sex-specific DNAm in the placenta affects genes that are fundamental to the placenta’s function.

A unique feature of our analysis was the exploration of placenta samples from earlier time points in pregnancy. Though these datasets were not conducive to genome-wide discovery owing to their small sample sizes, they did provide an ability to infer which CpG sites and regions identified as differentially methylated in full term samples displayed similar patterns of DNAm throughout the gestational period. We identified 194 CpG sites that met p-value thresholds of differential DNAm, with consistent direction of effect, in all four analyzed datasets (1st Trimester through full term). A protein-protein interaction analysis of the genes nearest to these sites implicated numerous central nervous system-related pathways (neurogenesis, axonogenesis, etc), as well as similar keratinization and peptide cross-linking pathways as seen in the full term analysis. As placenta datasets continue to be generated, it will be increasingly possible to better characterize persistently differentially methylated sites by fetal sex, to a greater degree possible than this study. Identification of these specific sites (and the genes and pathways they implicate) will be necessary to gain a greater understanding of the placenta’s role in “programming” sex-discordant phenotypes throughout the life course^3–5^.

At the region level, the two top-ranked DMRs from the full term analysis showed robust evidence for persistence across the gestational period. The top-ranked DMR, located in the CpG-island promoter of *ZNF175*, is characterized by a 17% hypermethylation in male samples relative to female, implying greater *ZNF175* gene expression in female samples. *ZNF175* regulates the expression of several chemokine receptors, and has been previously demonstrated to be up-regulated in pre-eclamptic vs. healthy placentas^23,24^, though these studies examined the maternal portion of the tissue. Future studies should aim to dissect the potential role of *ZNF175* promoter DNAm in fetal placenta in pre-eclampsia, considering both the male hypermethylated DMR described in this study, as well as the interesting intersection of fetal sex and gestational age in pre-eclampsia prevalence. Specifically, a recent large meta-analysis examining fetal sex and various birth outcomes found female fetuses to be associated with preterm pre-eclampsia, but male fetuses to be associated with term and postterm pre-eclampsia^44^.

The second-ranked DMR at the *ZNF300* promoter also displayed male hypermethylation to a similar degree (17%), along with strong evidence for the persistence of this DNAm difference across the gestational period. *ZNF300* is a transcriptional repressor with multiple reported targets^45^. It has been linked to immune cell differentiation^46,47^ and tumorigenesis^48,49^, furthering the longstanding narrative noting the similarity in placenta cells and cancer cells with respect to their invasive properties^50^. Along these lines, we found through incorporation of the Yuan et al. placenta cell type data^35^ that fetal sex differences in *ZNF300* promoter DNAm are driven by differences seen in the trophoblast cell type, which is primarily responsible for placental invasion^51^.

In placenta specifically, it has been previously demonstrated that samples collected from twins discordant for intrauterine growth restriction (IUGR) showed significant hypermethylation at the *ZNF300* promoter in the IUGR samples^30^. Our companion paper expands this story further through integration of imputed genotyping and placenta morphology measurements with comprehensive DNAm profiling (whole genome bisulfite sequencing, WGBS)^52^. Specifically, this study also demonstrates the existence of hypermethylation in male placentas at the *ZNF300* promoter, but further shows that these DNAm signatures mediate *cis* and *trans* (X chromosome) effects of genetic variants on placenta area. Collectively, these findings demonstrate robust evidence for a fetal sex DMR in the *ZNF300* promoter that is present across pregnancy and driven by trophoblast cell types, and provide a molecular mechanism for the governance of placenta size by fetal genetic variation. Future (*in vitro*) work should aim to further investigate the directionality of these phenomena in trophoblast cells specifically, to understand if CpG island promoter hypermethylation leads to reduced *ZNF300* expression (as expected canonically) and the relationship between *ZNF300* expression and cell proliferation.

There are several limitations of our approach that need to be recognized. First, we strictly leveraged publicly available data, which have limited annotation. In some cases, we were able to perform quality control or sanity checks to address these shortcomings. For example we used SNP probes on the Illumina 450K Array to identify and remove duplicate samples that may have been submitted publicly more than once, and we used a gestational age prediction algorithm to confirm that our sample classification was consistent with that guided by available metadata. But in some cases these checks were not possible, such as in selecting samples from normal/typical pregnancies, which was done solely with the provided metadata. Another limitation was the use of datasets that used the 450K array, as newer technologies to query DNAm now exist. Our companion paper^52^ describes an interrogation of sex-related DNAm patterns via WGBS, a comprehensive query of the methylome, but it is currently not feasible to perform these measurements on a large sample size. A better balance of methylome coverage and sample throughput is struck with the next iteration of the 450K Array, the Methylation EPIC array. Finally, our study utilized bulk placenta samples, which are a mixture of placenta cell types. We did perform several post hoc analyses to disentangle the role of different cell types in our findings. However, future work should aim to interrogate fetal sex DNAm differences directly in individual placenta cell populations to better understand, globally, the extent to which different cell types drive sex-related DNAm differences in the placenta.

Overall, future work should seek to better understand the consequence of fetal sex DNAm differences by establishing connections with additional molecular, phenotypic, and morphological endpoints. For example, paired placenta DNAm and gene expression or simply better integration of these two data types can be used to translate DNAm differences into gene expression changes, particularly in genomic regions outside of CpG island promoters and gene bodies where expectations of these relationships are less well known. In addition, placenta DNAm studies of pregnancy outcomes should be fetal sex-stratified or evaluate an interaction with fetal sex, given the significant differences by fetal sex identified in this study. In a similar vein, studies that collect placenta samples should aim to follow-up subjects throughout childhood and into adulthood whenever possible, to better quantify the potential for fetal sex DNAm differences to “program” sex-discordant health outcomes. Finally, as in our companion paper, future placenta studies should aim to collect placental morphology phenotypes in order to relate observed DNAm changes to placenta size, shape, and structure.

In conclusion, this study presents profound differences in DNAm from full term placentas according to fetal sex at the site and region level. Many of these differences, including strong hypermethylated signals in male samples relative to females at the *ZNF175* and *ZNF300* promoters, show strong evidence for their existence throughout pregnancy. Further characterization of these DNAm signatures will enable improved understanding of sex-specificity in placenta structure and function, gestational environments, and health outcomes across the life course.

## Methods

### Study selection

We searched the Gene Expression Omnibus (GEO) database on July 19th, 2018 for Illumina 450K microarray data sets containing human placental tissue samples by applying the following search terms: “placental methylation” or “placenta methylation” and limiting to platform types of “methylation profiling by array”, “methylation profiling by genome tiling array”, and “methylation profiling by SNP array”. We manually reviewed the resulting 78 studies and removed any that did not use the 450K array or did not make the raw .idat files available. We also restricted our study samples to only those labeled as phenotypically normal or controls by the metadata available in GEO. To assign samples within each study to gestational age (GA) categories of full term, first trimester, second trimester, and third trimester, we used the gestational age variable included in the sample metadata in GEO. Specifically, the first trimester group was defined as GA < 14 weeks, the second trimester as 14 δ GA < 28 weeks, the third trimester as 28 δ GA < 37 weeks, and full term as GA > 37 weeks. If a GA variable was not available, we consulted the manuscript associated with the GEO dataset. A total of 14 GEO datasets comprising 783 samples were included in the analysis, including 637 full term, 33 1st trimester, 72 2nd trimester, and 41 3rd trimester samples (**Supplementary Table 1**). We predicted the gestational age of these samples using the *predictAge()* function from the planet R package^53^ to verify that our sample groups captured their intended points during the gestation period^54^.

### Array pre-processing, quality control, and batch effect correction

All analyses were performed using R (v4.0.3) and the *minfi* R package (v1.36)^55^ unless otherwise stated. We theorized that some of the placenta samples in the public domain could be duplicates. Hence, from the samples that passed the inclusion criteria, duplicate pairs of samples were identified using the 65 single nucleotide polymorphisms (SNP) probes measured on the 450K array. The sample from the identified pair with the greater number of probes defined as detection p-value failures (p > 0.01) was removed. The *preprocessNoob()* function^56,57^ was used for background correction and dye-bias equalization. Samples that had low overall intensity (median unmethylated or methylated signal < 11) or had a detection p-value > 0.01 in more than 1% of probes or probes that had a detection p-value > 0.01 in more than 10% of samples were removed. Samples were removed if the reported sex did not match predicted sex generated by the *getSex()* function in *minfi*. We removed probes mapping to sex chromosomes, and also removed autosomal-targeting probes that ambiguously map to sex chromosome locations according to Chen et al.^58^. After these steps the dataset for the full term samples consisted of 445,278 probes and 532 samples. Finally, quantile normalization was performed, and surrogate variables were estimated using the sva R package^59^ on the resulting M-values. SVs have been shown to capture differences related to batch effects and cell type proportions across samples in a wide variety of simulated settings^60^, and to remove the effects of unwanted sources of technical and biological variation^59^.

We repeated these QC and batch effect correction steps in each trimester dataset individually. For the 1st trimester dataset, these steps resulted in 444,948 probes and 22 samples available for downstream analysis. For the 2nd trimester, it was 445,331 probes and 59 samples and for the 3rd trimester it was 445,220 probes and 37 samples.

### Identification of differentially methylated CpG sites and regions

To identify differentially methylated individual CpG sites using the *limma* R package^61^, we modeled M-values as function of fetal sex and the estimated SVs and defined genome-wide significant probes as those with p-values < 1E-8. The *bumphunter()* function^62^ was used to identify differentially methylated regions (DMRs) using the same model. We repeated these same association testing procedures in each trimester individually. Because these datasets were not powered for genome-wide discovery, we did not impose strict statistical significance cutoffs for these analyses. Instead, we looked for evidence of replication of differentially DNAm identified in the full term samples, to quantify the extent to which these differences persisted across the gestational period. We considered a early term probe to replicate the full term result if the fetal sex analysis showed a p-value < 0.05 and a consistent direction of effect with that observed in the full term samples.

### Gene enrichment and pathway analysis

After identifying differentially methylated CpG sites and DMRs, we sought to characterize biological pathways enriched in our top findings. All probes reaching a significance threshold of 1E-8 were evaluated for enrichment in Gene Ontology (GO) pathways using the *gometh()* and *topGSA()* functions in the R package *missMethyl*. STRING protein-protein interaction network analyses were also performed after mapping significant CpG sites to their nearest genes^63^.

### Comparison of results to previous DNAm studies of sex

Significant CpG sites discovered via our analysis were compared to significant CpG sites found in previous human sex-related DNAm studies to determine how many sites had already been identified in the literature. These previous studies analyzed placenta^41,64^ as well as cord blood^65,66^, peripheral blood^66–69^, saliva^70^, prefrontal cortex^71,72^, pancreas^73^, and fetal cortex^74^.

### Cell type heterogeneity analyses

To understand the extent to which our fetal sex differentially methylated sites and regions were driven by cell type heterogeneity, we compared our results to those from a recent study by Yuan et al. examining placenta cell type-specific DNAm patterns^35^. Specifically, we quantified the degree of overlap in differentially methylated sites from our full term analysis with probes identified in the Yuan et al. study as distinguishing cell type at full term, and conducted GO pathway analyses in groups of probes that did and did not overlap with this list separately. We also determined which of the identified DMRs from the full term analysis contained probes from this list, downloaded the data from the Yuan et al. study (GEO ID: GSE159526) and plotted the cell type specific DNAm profiles in these same regions to determine which cell type(s) were driving the DMR result.

## Supporting information

Description of Supplementary Data

## Data Availability

All placenta datasets used for association testing in this study are available in GEO and listed in **Supplementary Table 1**. Cell type specific placenta DNAm data described in Yuan et al.^35^ were also downloaded from GEO (GSE159526).

## Code Availability

Scripts for analyses conducted in this study are available at https://github.com/sandrews5/PlacentaDNAm_FetalSex

## Declarations

## Author’s Contributions

SVA conceived the study, performed the data analysis, and wrote the paper. IJY performed the data analysis and wrote the paper. KF performed the data analysis and wrote the paper. TO wrote the paper. MS conceived the study and wrote the paper. All authors contributed to interpretation of results and edited and reviewed the manuscript.

## Correspondence

Correspondence to Marina Sirota

