## Supplementary Information for "Large-scale placenta DNA methylation mega-analysis reveals fetal sex-specific differentially methylated CpG sites and regions"

**Supplementary Table 1: Datasets included in this study.** Included Full Term Analysis: Y/N variable indicating if samples from dataset were used in full term analysis. Inclusion Criteria Full Term Analysis: criteria to select samples for full term analysis. Included Early Term Analysis: Y/N variable indicating if samples from dataset were used in any of the trimester specific analyses. Inclusion Criteria Early Term Analysis: criteria to select samples for inclusion in trimester specific cohorts. N_Term/1stTerm/2ndTerm/3rdTerm: Number of samples (pre QC) included from dataset in each analysis subgroup.

| GEOID | Included Full Term Analysis | Inclusion Criteria Full Term Analysis | Included Early Term Analysis | Inclusion Criteria Early Term Analysis | N term | N 1stTerm | N 2ndTerm | N 3rdTerm |
|---|---|---|---|---|---|---|---|---|
| GSE108567 | Y | Via Gestational Age | Y | Via Gestational Age | 45 | NA | NA | 14 |
| GSE100197 | Y | Term control | Y | Preterm control | 19 | NA | 2 | 22 |
| GSE98224 | Y | Term control, AGA | Y | Preterm, control, AGA | 9 | NA | NA | 5 |
| GSE106089 | N | NA | Y | All | NA | NA | 46 | NA |
| GSE103413 | N | NA | Y | Chorionic villus | NA | NA | 2 | NA |
| GSE98938 | Y | chorion, trophoblast, villi | Y | chorion, villi, trophoblast | 6 | NA | 6 | NA |
| GSE93208 | N | NA | Y | All | NA | 19 | NA | NA |
| GSE71719 | Y | Bisulfite converted | N | NA | 23 | NA | NA | NA |
| GSE71678 | Y | All | N | NA | 323 | NA | NA | NA |
| GSE75248 | Y | AGA samples | N | NA | 174 | NA | NA | NA |
| GSE75196 | Y | Healthy | N | NA | 16 | NA | NA | NA |
| GSE69502 | N | NA | Y | Chorionic Villi, Controls | NA | NA | 16 | NA |
| GSE74738 | Y | Controls | N | NA | 22 | NA | NA | NA |
| GSE66210 | N | NA | N | Chorionic villus, normal | NA | 14 | NA | NA |
| | | | | **SUM** | 637 | 33 | 72 | 41 |

### *Supplementary Figures*

**Supplementary Figure 1: PCA plot of 3rd Trimester placenta samples before and after batch effect correction.** Top triangle shows principal components generated from methylation M-values at all autosomal probes passing QC prior to quantile normalization or surrogate variable analysis. Bottom triangle shows residuals of quantile normalized autosomal M-values regressed onto estimated surrogate variables.

**Supplementary Figure 2: PCA plot of 2nd Trimester placenta samples before and after batch effect correction.** Top triangle shows principal components generated from methylation M-values at all autosomal probes passing QC prior to quantile normalization or surrogate variable analysis. Bottom triangle shows residuals of quantile normalized autosomal M-values regressed onto estimated surrogate variables.

**Supplementary Figure 3: PCA plot of 1st Trimester placenta samples before and after batch effect correction.** Top triangle shows principal components generated from methylation M-values at all autosomal probes passing QC prior to quantile normalization or surrogate variable analysis. Bottom triangle shows residuals of quantile normalized autosomal M-values regressed onto estimated surrogate variables.

**Supplementary Figure 4: Distribution of absolute value of mean differences in sites that do discriminate placenta cell types (gray, n = 1,285) and sites that do not (orange, n = 3927), among the genome-wide significant sites from the full term analysis.** Cell type discriminating probes determined by Yuan et al.

**Supplementary Figure 5: Manhattan plots showing single site association results for each early term analysis.**

**Supplementary Figure 6: Volcano plot for full term DMR analysis.** The x-axis measures the average methylation difference between male and female fetuses, and the y-axis measures the size of the DMR in base pairs. The blue points represent DMRs considered significant (fwerArea < 0.1)

**Supplementary Figures 7 - 14: Additional plots depicting significant DMRs from full term analysis across all gestational periods.** Remaining DMRs exceeding fwerArea threshold of 0.1 as seen in Table 4 (excluding top-ranked DMR in ZNF175 promoter, see Figure 4). In each figure, DMRs are plotted as percent methylation as a function of genomic position. Dots indicate samples and solid lines indicate smooth lines through male sample values (blue) and female sample values (gold). Actual identified DMR region indicated via dashed lines. **a** Full Term Samples. **b** 1st Trimester Samples **c** 2nd Trimester Samples **d** 3rd Trimester Samples.

**Supplementary Figures 15 - 18: Exploration of cell type heterogeneity in additional DMRs.** Remaining DMRs containing cell type discriminating probes as determined by Yuan et al (see Table 4). DMRs are plotted as percent methylation as a function of genomic position. Dots indicate samples and solid lines indicate smooth lines through male sample values (blue) and female sample values (gold). Actual identified DMR region indicated via dashed lines. **a** DMR identified in full term analysis, plotted using full term samples from present study. **b - d** The same region, but plotted using samples from different datasets. **b** Term samples and individual placenta cell types from Yuan et al. **c** 1st Trimester samples from present study. **d** 1st trimester samples and individual placenta cell types from Yuan et al. In **b** + **d**, fractions indicate the proportion of probes in the region that were annotated by Yuan et al. as distinguishing that cell type in full term and 1st trimester samples, respectively. Villi samples were not evaluated in this manner by Yuan et al. as this is the unsorted bulk tissue.

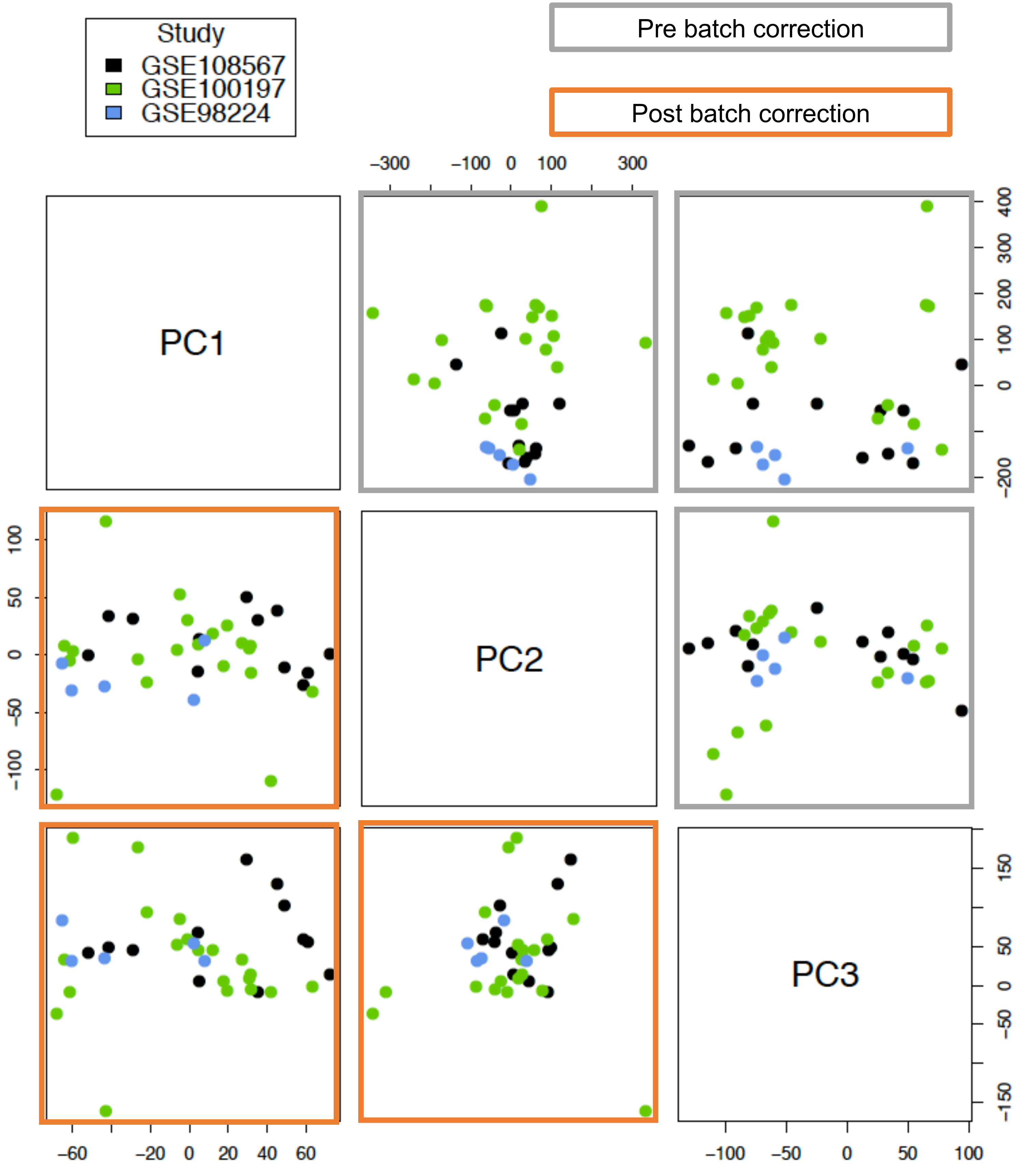
Study
GSE108567
GSE100197
GSE98224
Pre batch correction
Post batch correction
PC1
PC2
PC3
−300
−100
0
100
300
400
300
200
100
0
−200
100
50
0
−50
−100
150
50
0
−50
−150
−60
−20
0
20
40
60
−100
−50
0
50
100

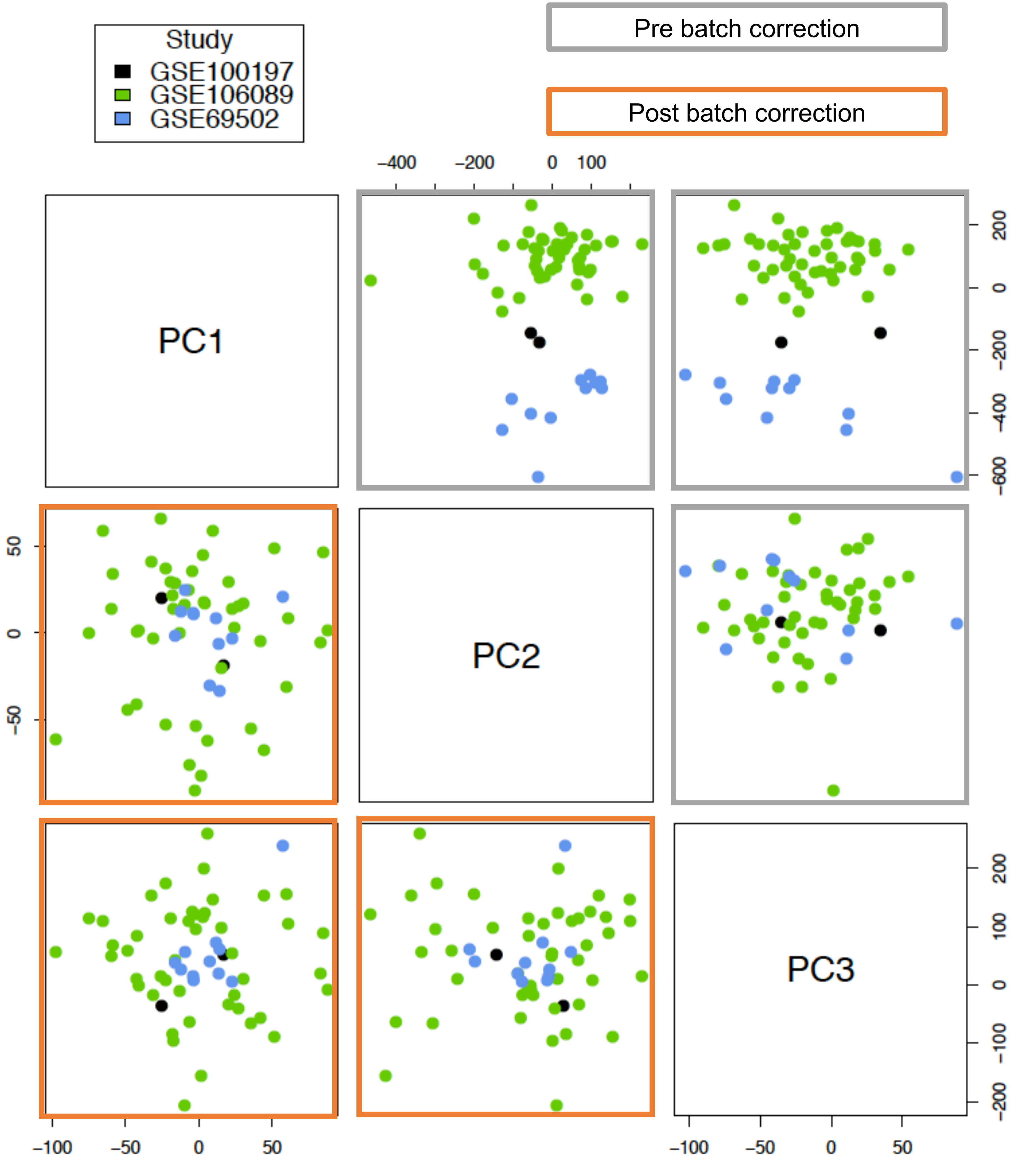

Study
GSE100197
GSE106089
GSE69502
Pre batch correction
Post batch correction
PC1
PC2
PC3
−400
−200
0
100
200
0
−200
−400
−600
50
0
−50
200
100
0
−100
−200
−100
−50
0
50
−100
−50
0
50

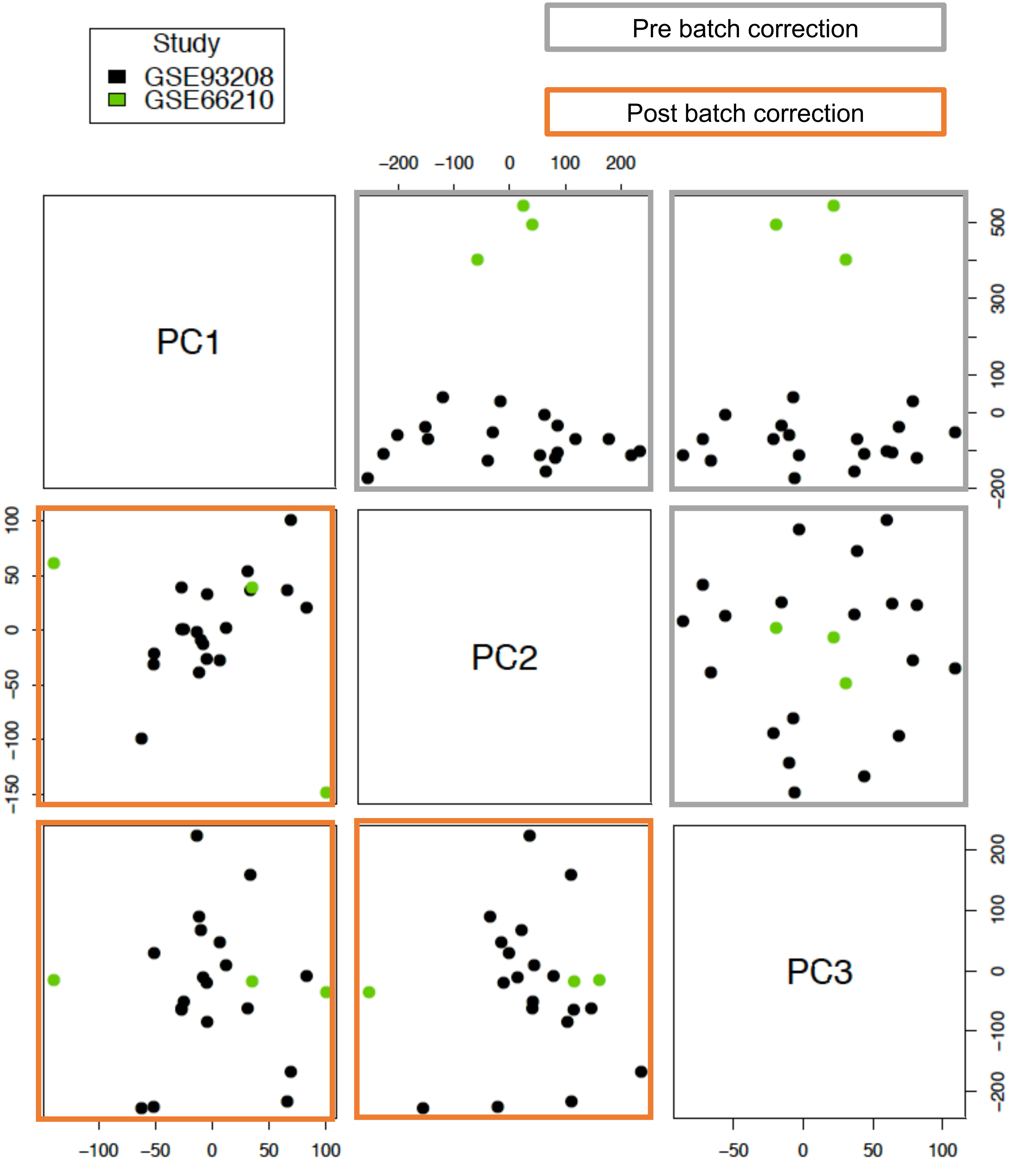
Study
GSE93208
GSE66210
Pre batch correction
Post batch correction
PC1
PC2
PC3
−200 −100 0 100 200
500 300 100 0 −200
100 50 0 −50 −100 −150
200 100 0 −100 −200
−100 −50 0 50 100
−50 0 50 100

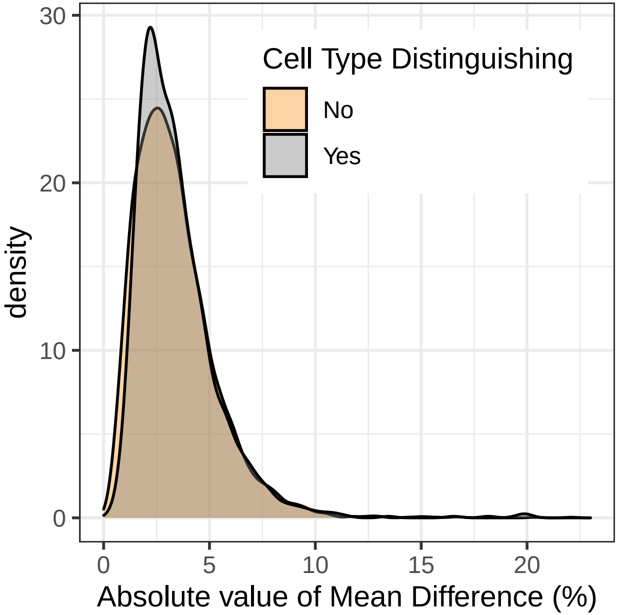

Cell Type Distinguishing
No
Yes
density
Absolute value of Mean Difference (%)
0
10
20
30
0
5
10
15
20

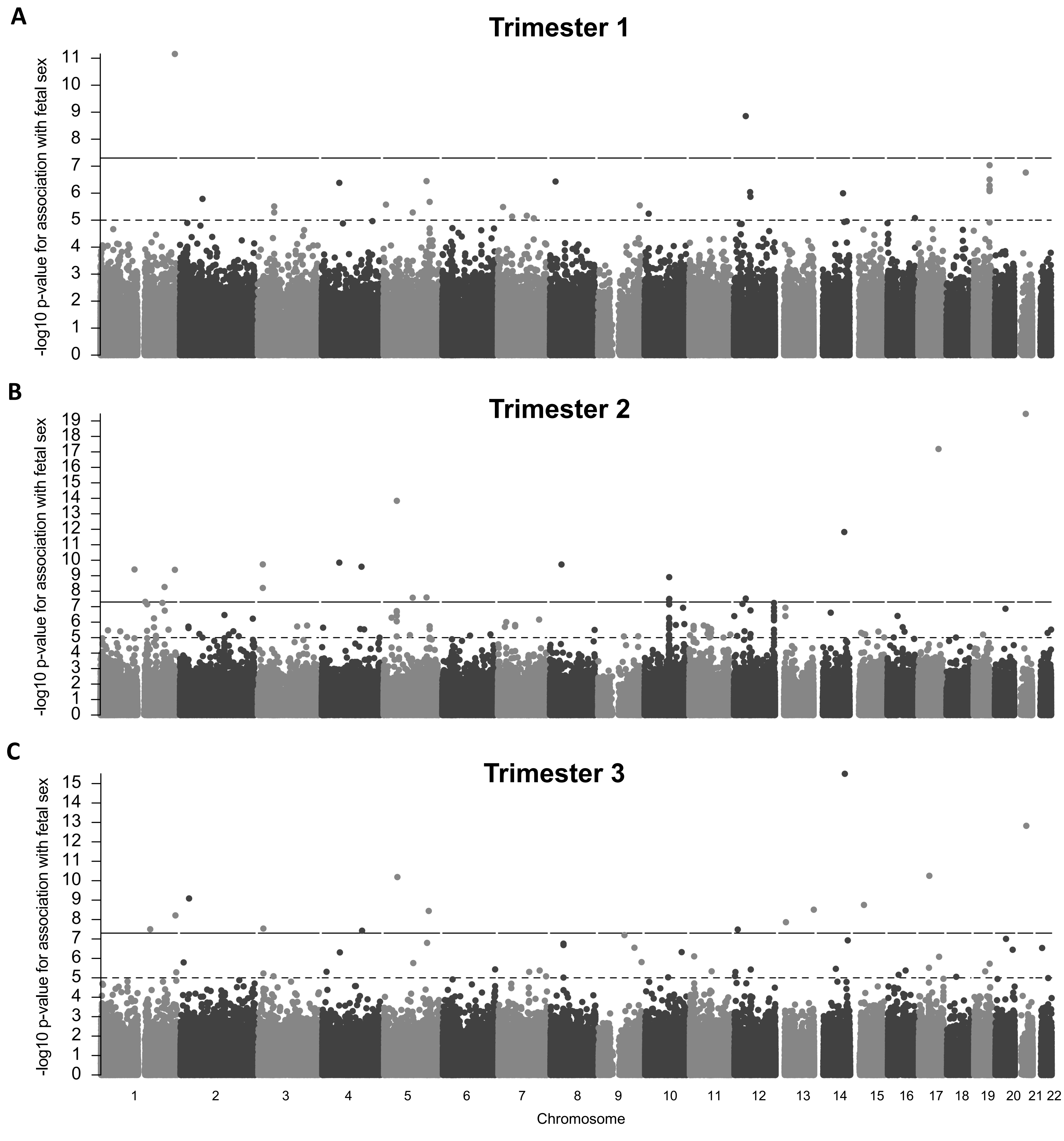
A
Trimester 1
B
Trimester 2
C
Trimester 3
-log10 p-value for association with fetal sex
Chromosome
1 2 3 4 5 6 7 8 9 10 11 12 13 14 15 16 17 18 19 20 21 22

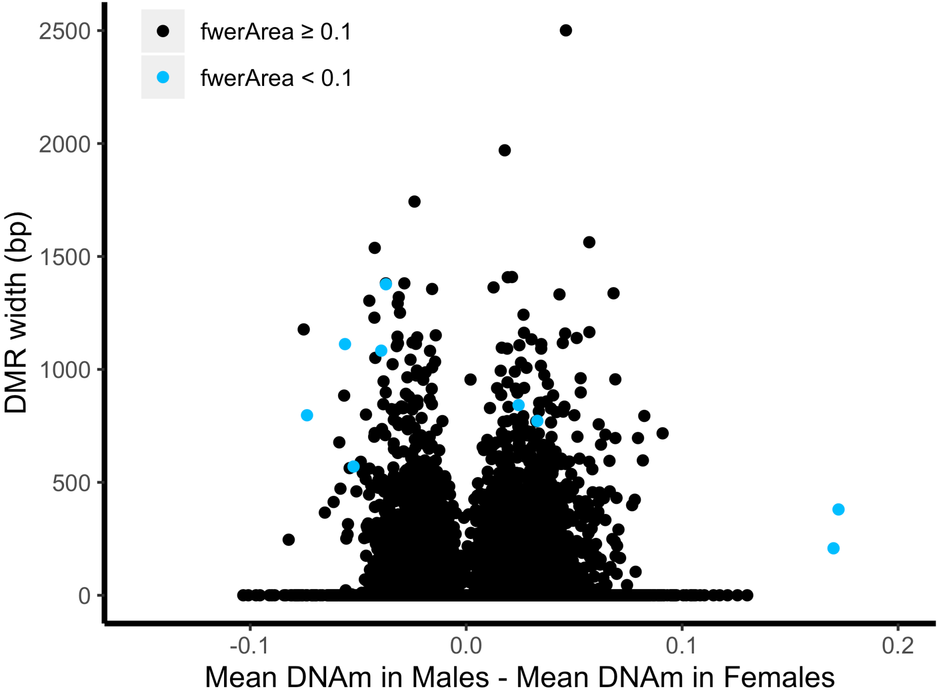

fwerArea ≥ 0.1
fwerArea < 0.1
DMR width (bp)
2500
2000
1500
1000
500
0
-0.1
0.0
0.1
0.2
Mean DNAm in Males - Mean DNAm in Females

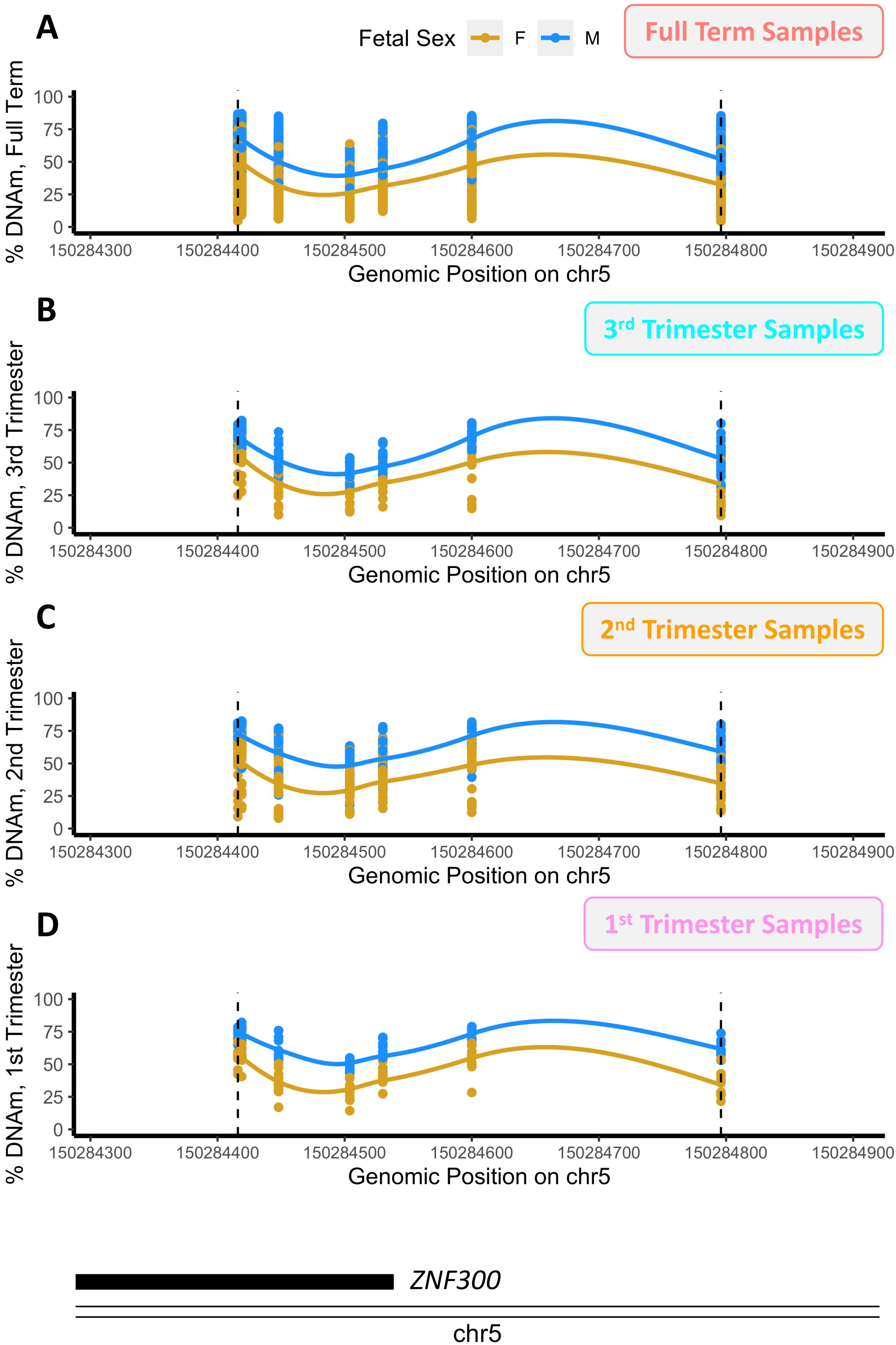

A
Fetal Sex
F
M
Full Term Samples
% DNAm, Full Term
Genomic Position on chr5
B
3rd Trimester Samples
% DNAm, 3rd Trimester
Genomic Position on chr5
C
2nd Trimester Samples
% DNAm, 2nd Trimester
Genomic Position on chr5
D
1st Trimester Samples
% DNAm, 1st Trimester
Genomic Position on chr5
100
75
50
25
0
150284300
150284400
150284500
150284600
150284700
150284800
150284900
ZNF300
chr5

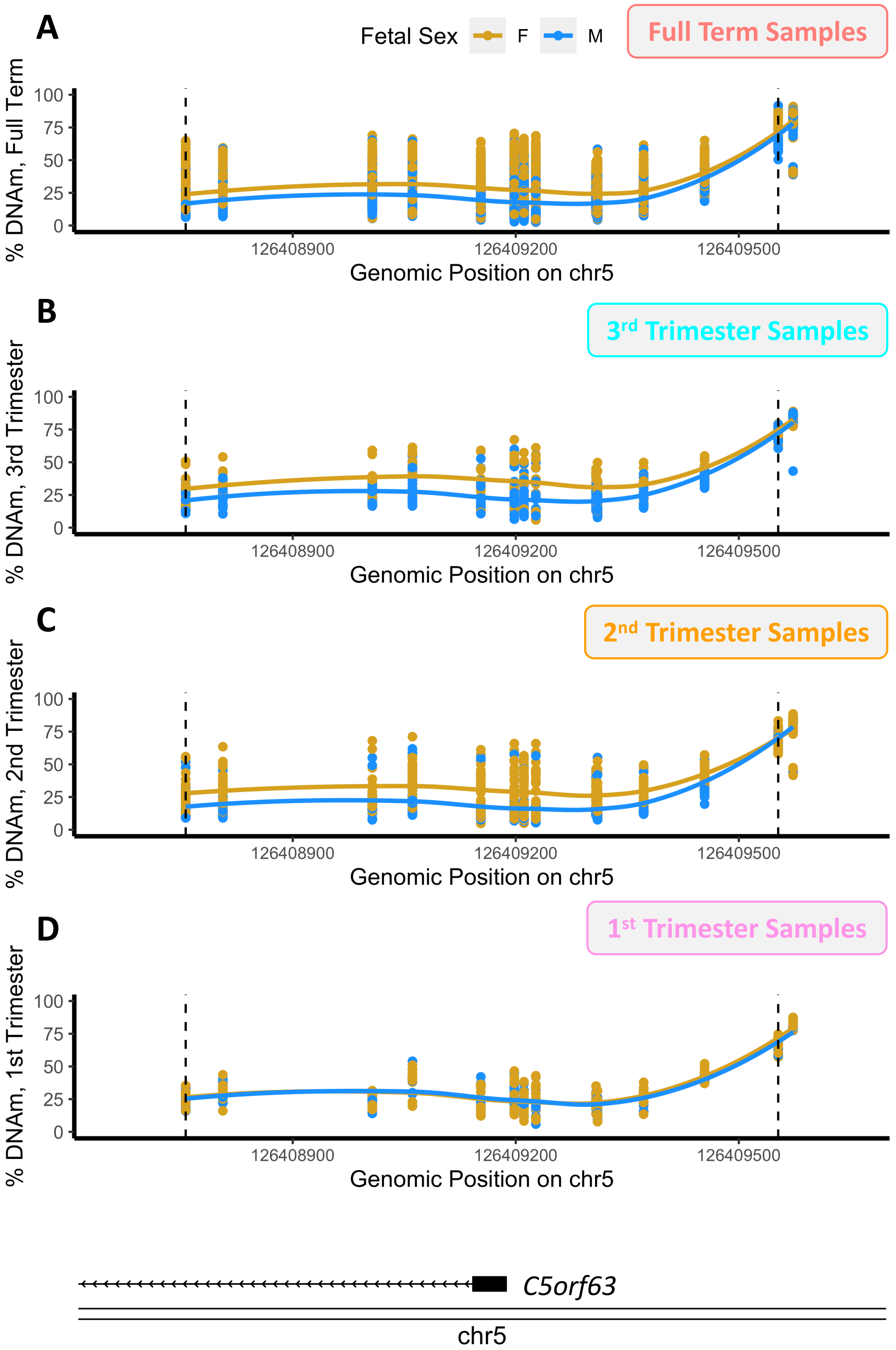

A
Fetal Sex
F
M
Full Term Samples
% DNAm, Full Term
100
75
50
25
0
126408900
126409200
126409500
Genomic Position on chr5
B
3rd Trimester Samples
% DNAm, 3rd Trimester
100
75
50
25
0
126408900
126409200
126409500
Genomic Position on chr5
C
2nd Trimester Samples
% DNAm, 2nd Trimester
100
75
50
25
0
126408900
126409200
126409500
Genomic Position on chr5
D
1st Trimester Samples
% DNAm, 1st Trimester
100
75
50
25
0
126408900
126409200
126409500
Genomic Position on chr5
C5orf63
chr5

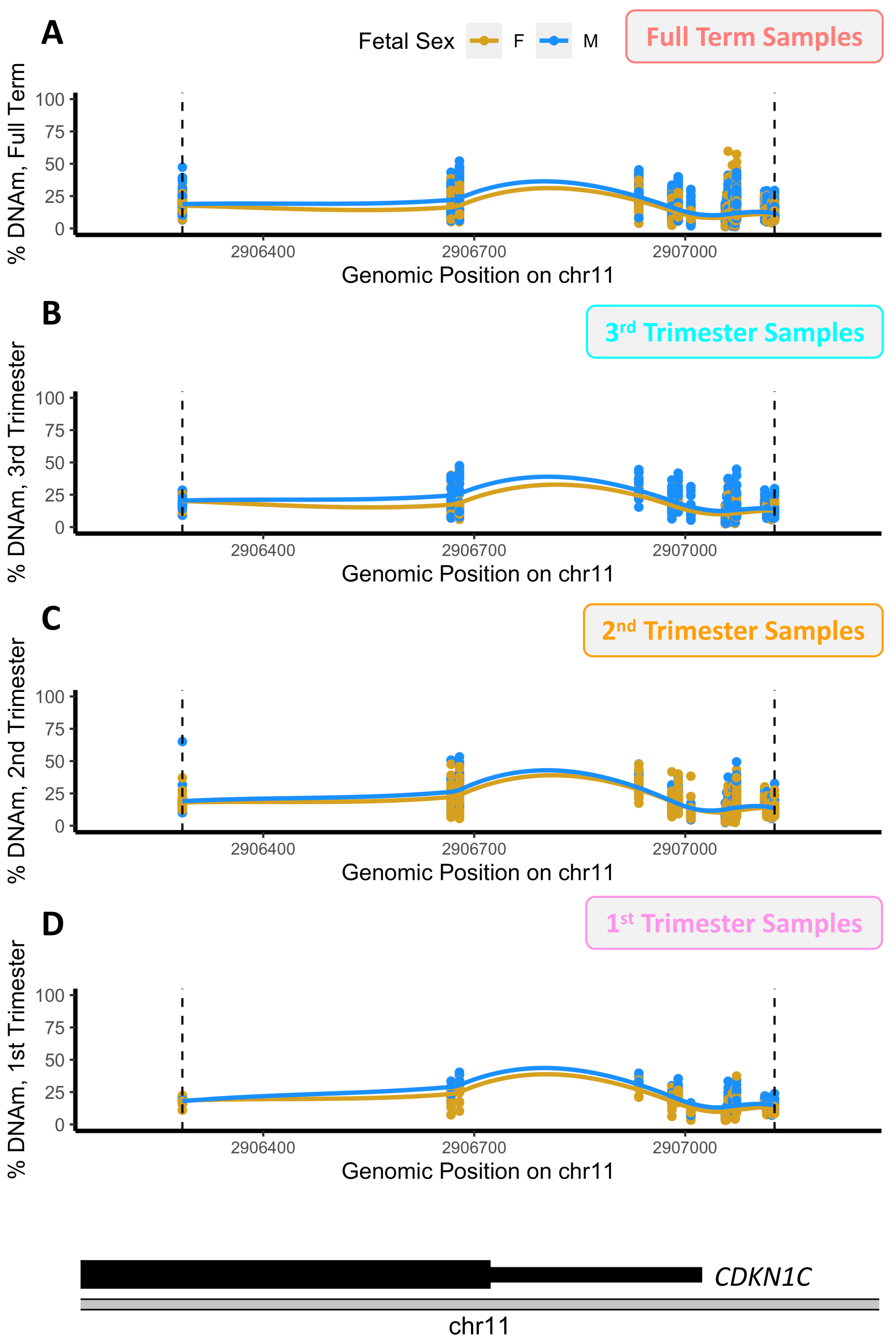

A
Fetal Sex
F
M
Full Term Samples
% DNAm, Full Term
100
75
50
25
0
2906400
2906700
2907000
Genomic Position on chr11
B
3rd Trimester Samples
% DNAm, 3rd Trimester
C
2nd Trimester Samples
% DNAm, 2nd Trimester
D
1st Trimester Samples
% DNAm, 1st Trimester
CDKN1C
chr11

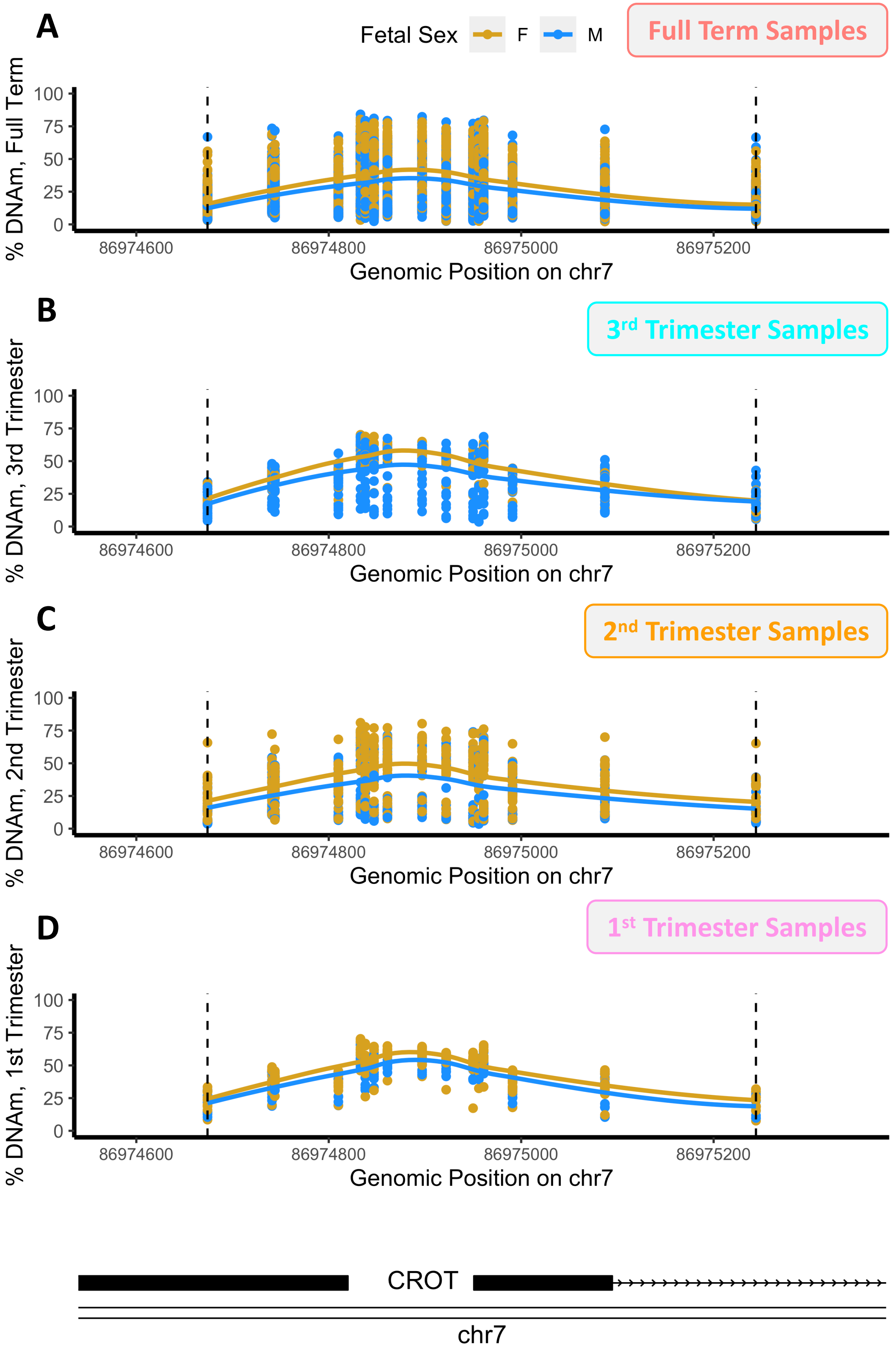

A
Fetal Sex
F
M
Full Term Samples
% DNAm, Full Term
100
75
50
25
0
86974600
86974800
86975000
86975200
Genomic Position on chr7
B
3rd Trimester Samples
% DNAm, 3rd Trimester
C
2nd Trimester Samples
% DNAm, 2nd Trimester
D
1st Trimester Samples
% DNAm, 1st Trimester
CROT
chr7

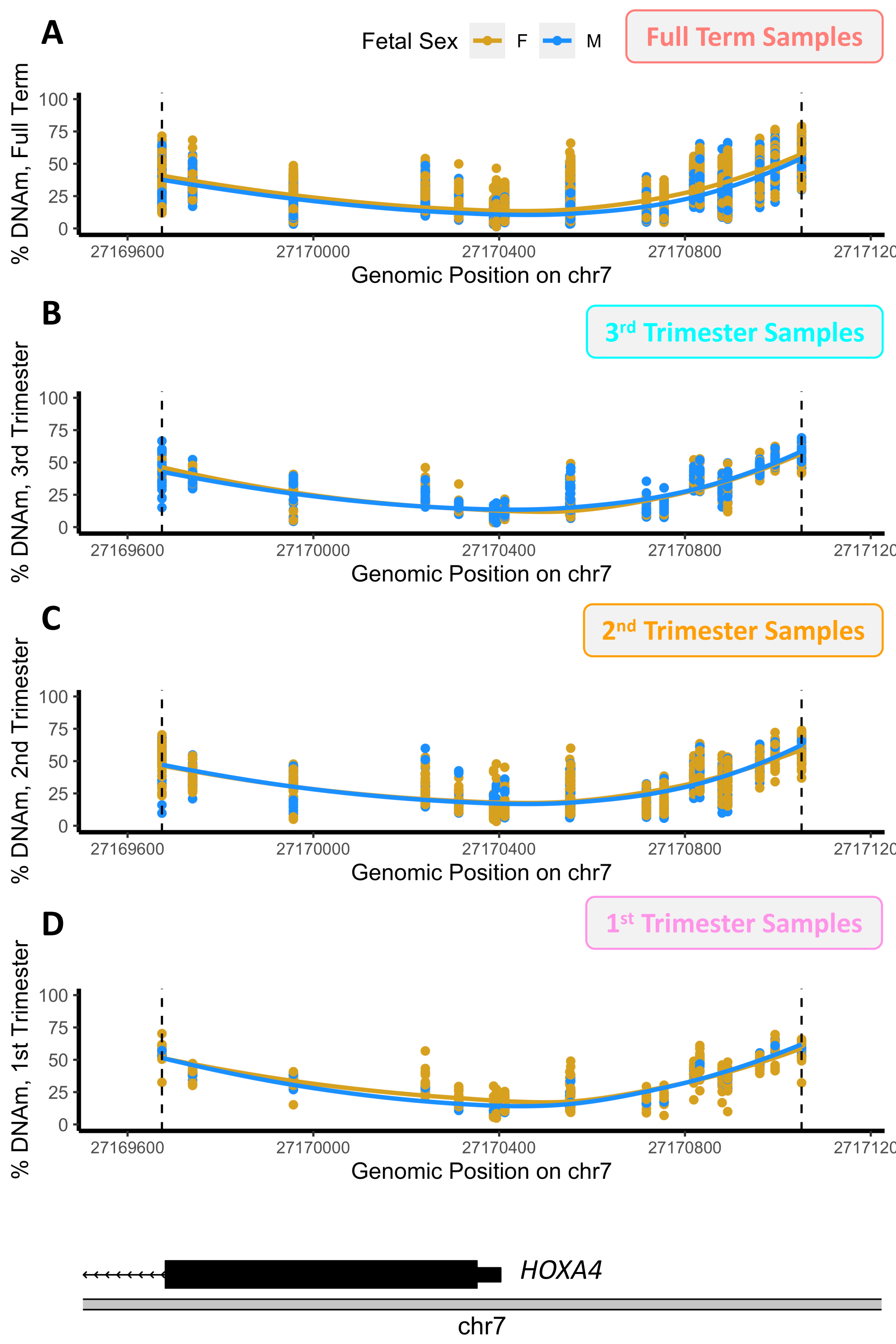

A
Fetal Sex
F
M
Full Term Samples
% DNAm, Full Term
100
75
50
25
0
27169600
27170000
27170400
27170800
Genomic Position on chr7
B
3rd Trimester Samples
% DNAm, 3rd Trimester
Genomic Position on chr7
C
2nd Trimester Samples
% DNAm, 2nd Trimester
Genomic Position on chr7
D
1st Trimester Samples
% DNAm, 1st Trimester
Genomic Position on chr7
HOXA4
chr7

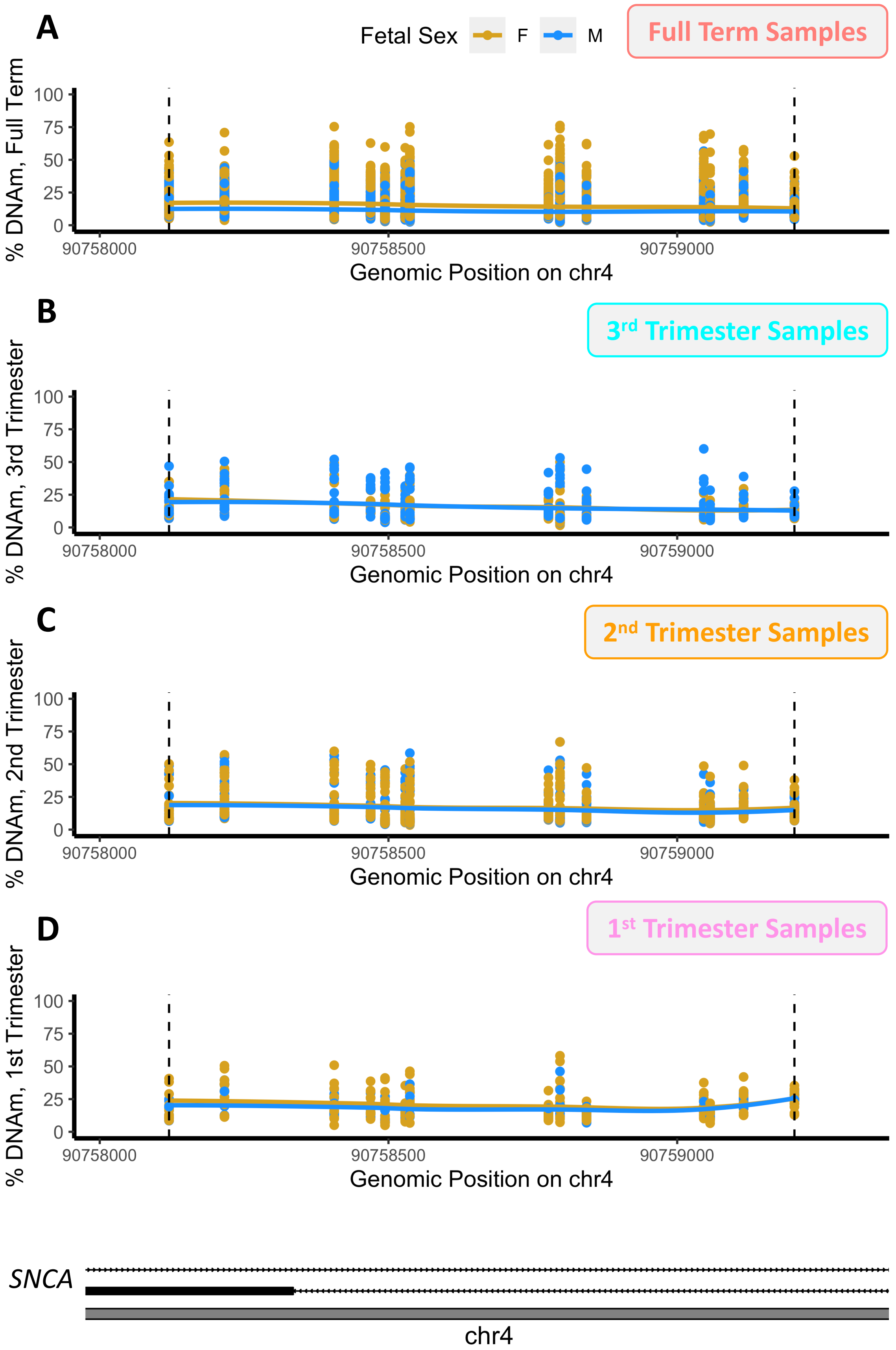

A
Fetal Sex
F
M
Full Term Samples
% DNAm, Full Term
Genomic Position on chr4
B
3rd Trimester Samples
% DNAm, 3rd Trimester
C
2nd Trimester Samples
% DNAm, 2nd Trimester
D
1st Trimester Samples
% DNAm, 1st Trimester
0
25
50
75
100
90758000
90758500
90759000
SNCA
chr4

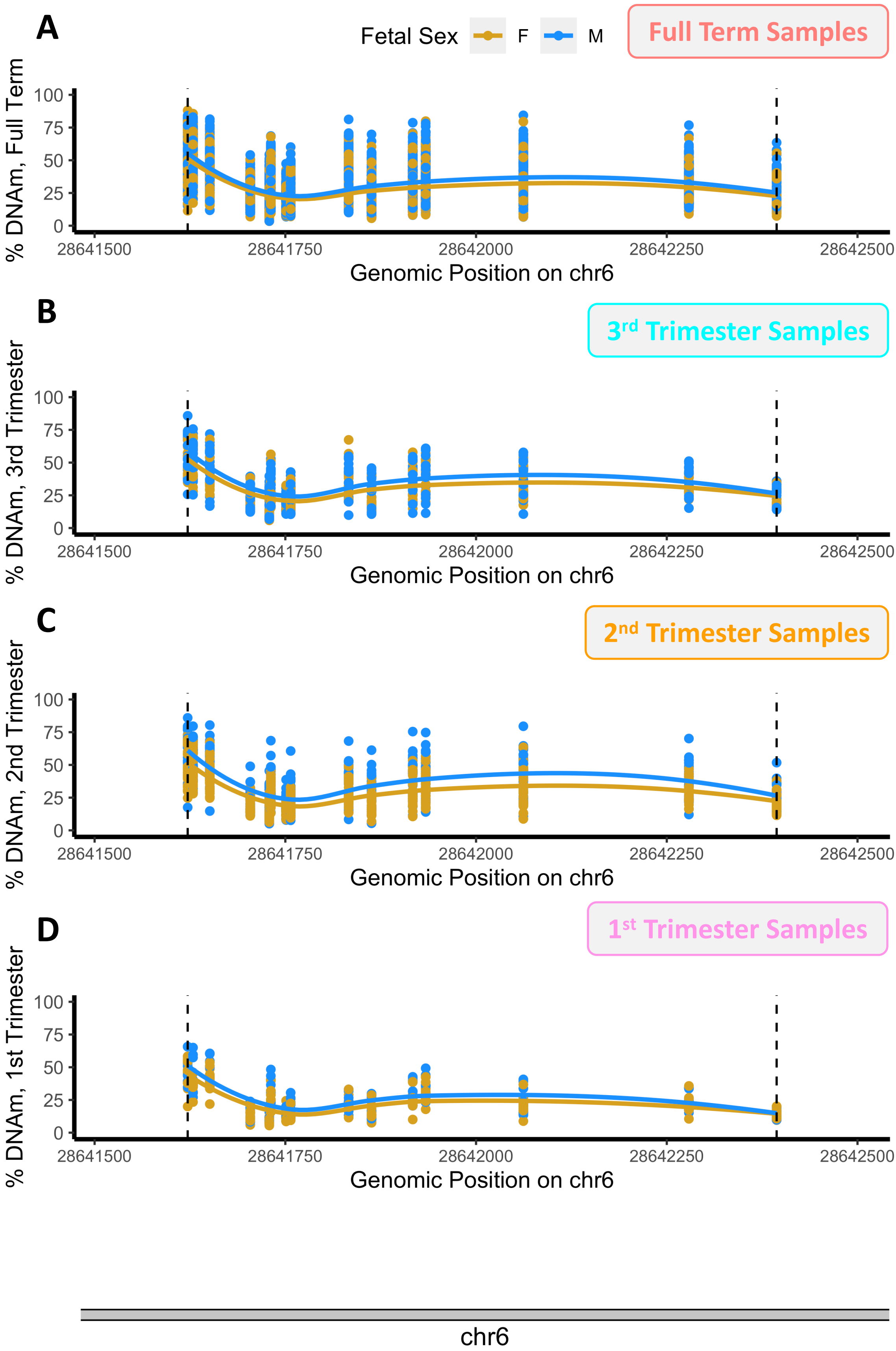

A
Fetal Sex
F
M
Full Term Samples
% DNAm, Full Term
100
75
50
25
0
28641500
28641750
28642000
28642250
28642500
Genomic Position on chr6
B
3rd Trimester Samples
% DNAm, 3rd Trimester
C
2nd Trimester Samples
% DNAm, 2nd Trimester
D
1st Trimester Samples
% DNAm, 1st Trimester
chr6

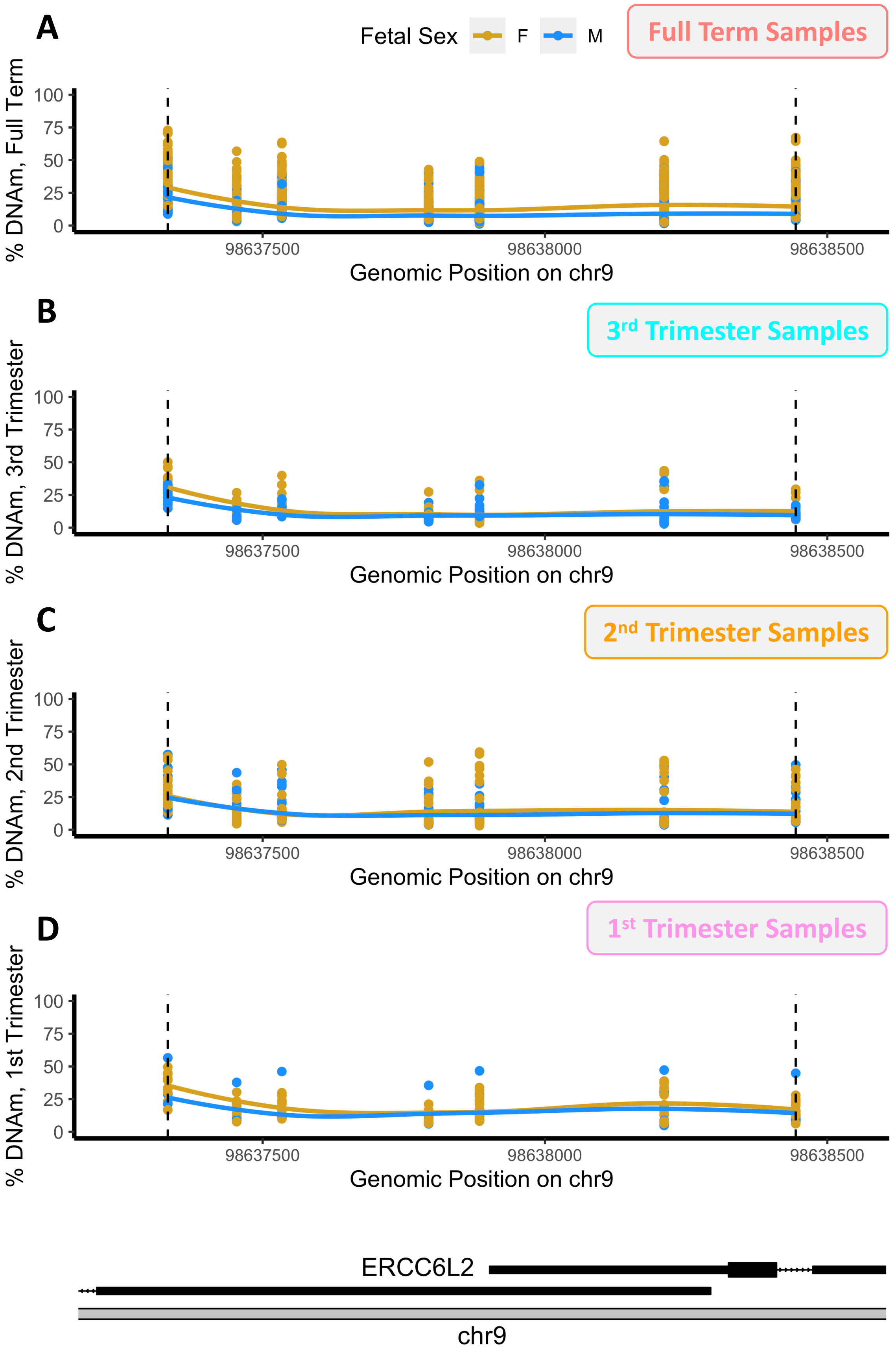
A
Fetal Sex
F
M
Full Term Samples
% DNAm, Full Term
100
75
50
25
0
98637500
98638000
98638500
Genomic Position on chr9
B
3rd Trimester Samples
% DNAm, 3rd Trimester
Genomic Position on chr9
C
2nd Trimester Samples
% DNAm, 2nd Trimester
Genomic Position on chr9
D
1st Trimester Samples
% DNAm, 1st Trimester
Genomic Position on chr9
ERCC6L2
chr9

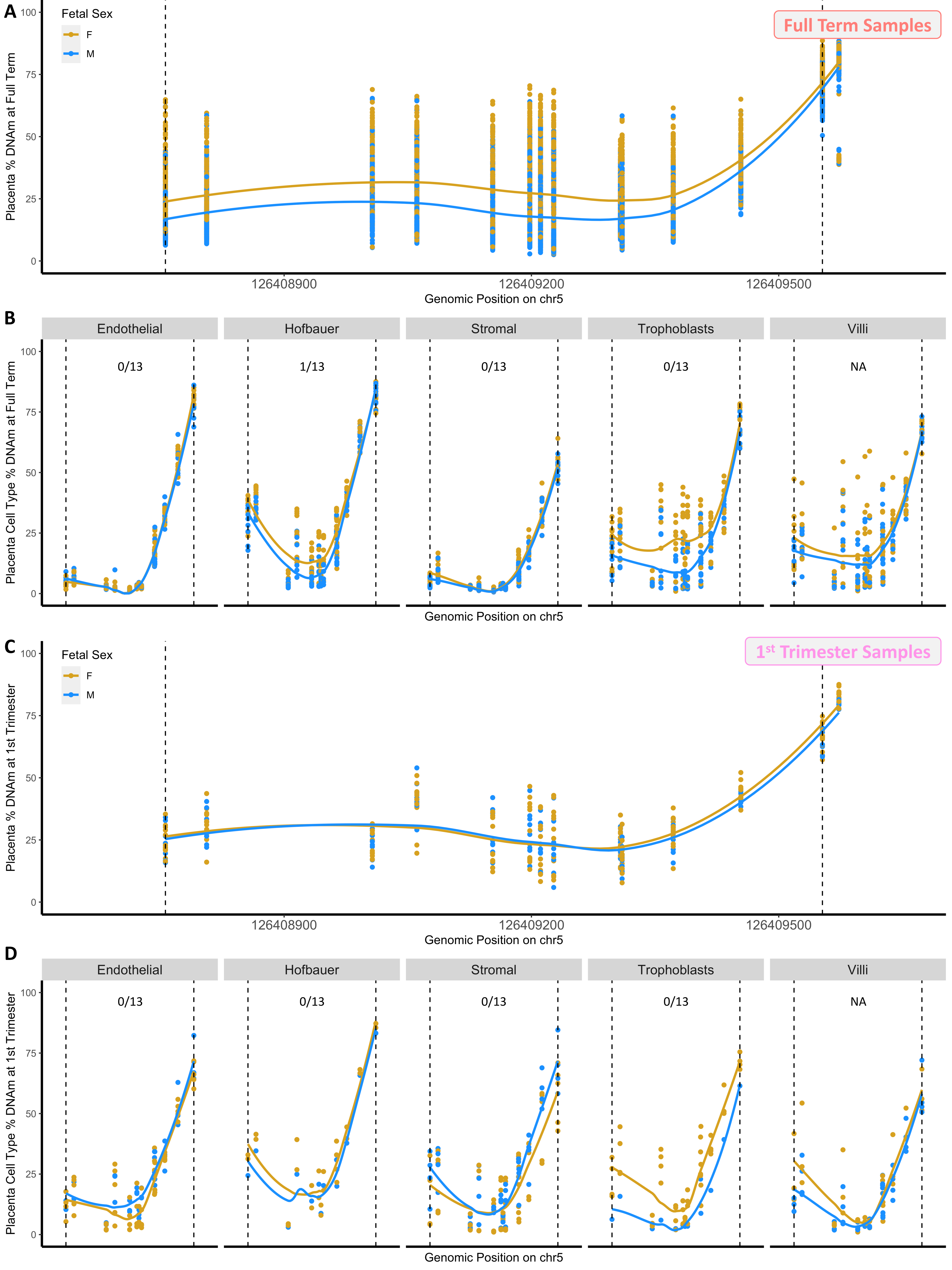
A
Fetal Sex
F
M
Full Term Samples
Placenta % DNAm at Full Term
100
75
50
25
0
126408900
126409200
126409500
Genomic Position on chr5
B
Endothelial
Hofbauer
Stromal
Trophoblasts
Villi
0/13
1/13
0/13
0/13
NA
Placenta Cell Type % DNAm at Full Term
Genomic Position on chr5
C
Fetal Sex
F
M
1st Trimester Samples
Placenta % DNAm at 1st Trimester
126408900
126409200
126409500
Genomic Position on chr5
D
Endothelial
Hofbauer
Stromal
Trophoblasts
Villi
0/13
0/13
0/13
0/13
NA
Placenta Cell Type % DNAm at 1st Trimester
Genomic Position on chr5

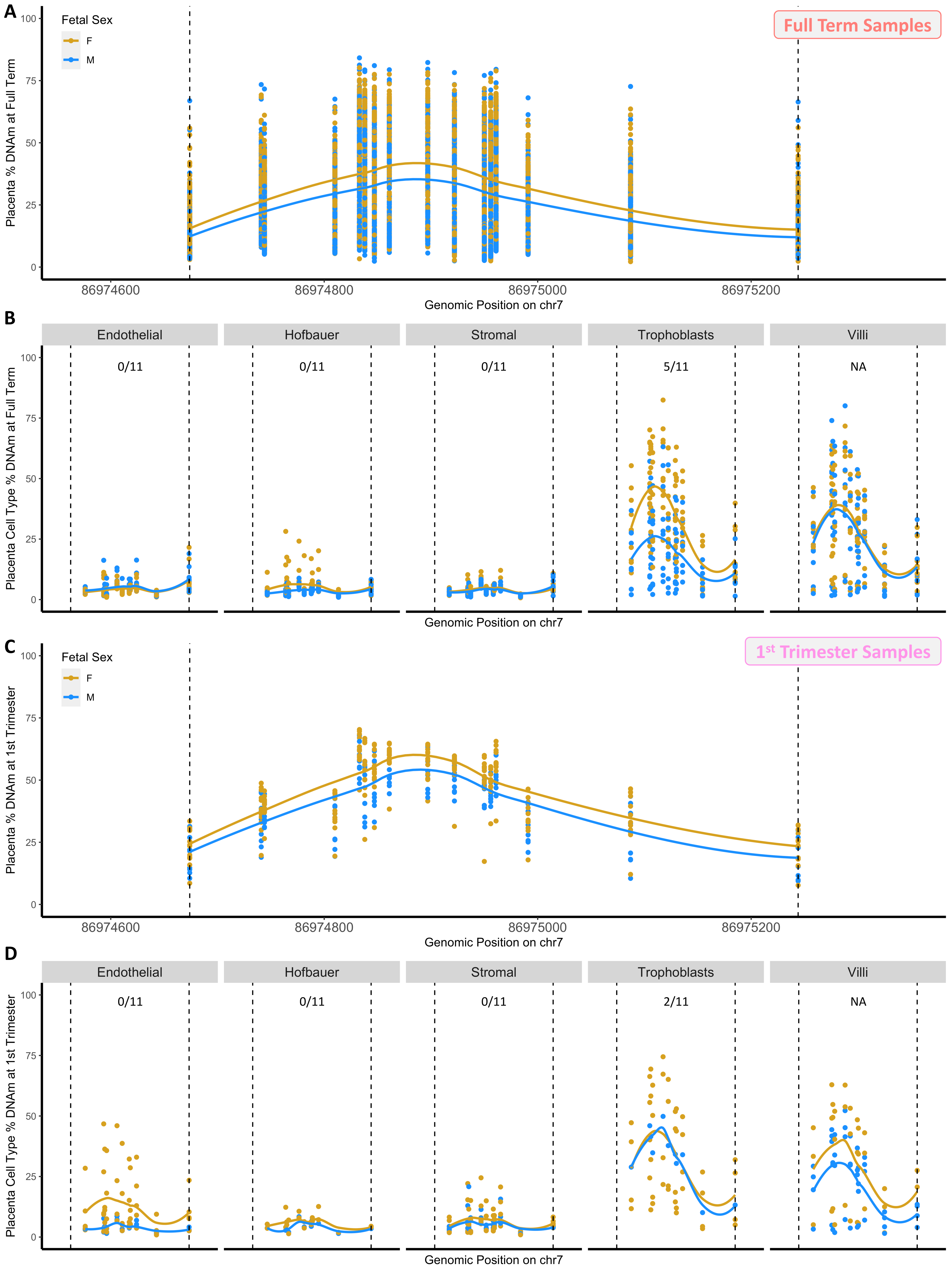
A
Full Term Samples
Fetal Sex
F
M
Placenta % DNAm at Full Term
Genomic Position on chr7
B
Endothelial
Hofbauer
Stromal
Trophoblasts
Villi
0/11
0/11
0/11
5/11
NA
Placenta Cell Type % DNAm at Full Term
Genomic Position on chr7
C
1st Trimester Samples
Fetal Sex
F
M
Placenta % DNAm at 1st Trimester
Genomic Position on chr7
D
Endothelial
Hofbauer
Stromal
Trophoblasts
Villi
0/11
0/11
0/11
2/11
NA
Placenta Cell Type % DNAm at 1st Trimester
Genomic Position on chr7

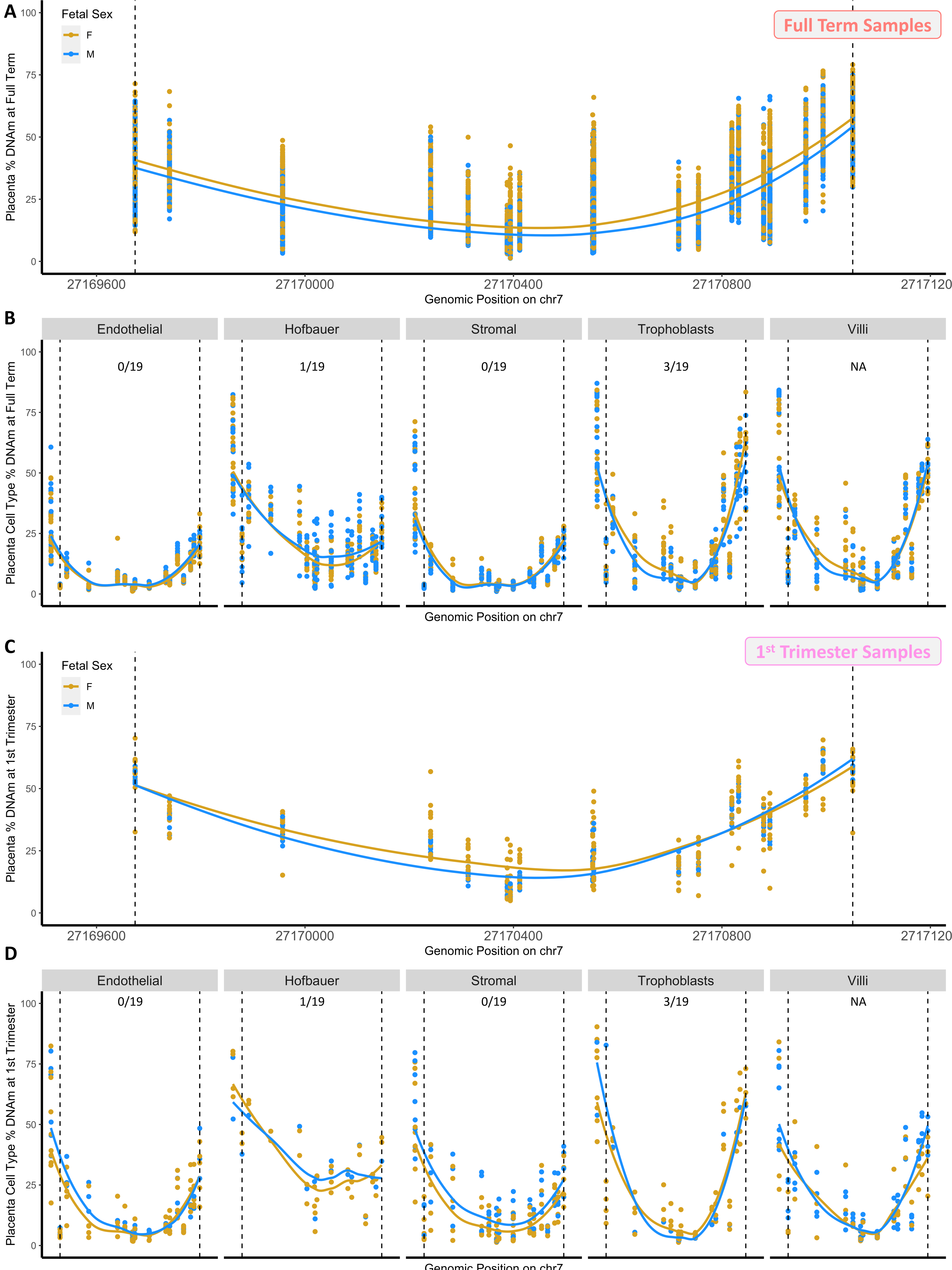
A
Full Term Samples
Fetal Sex
F
M
Placenta % DNAm at Full Term
Genomic Position on chr7
27169600
27170000
27170400
27170800
2717120
B
Endothelial
Hofbauer
Stromal
Trophoblasts
Villi
0/19
1/19
0/19
3/19
NA
Placenta Cell Type % DNAm at Full Term
Genomic Position on chr7
C
1st Trimester Samples
Fetal Sex
F
M
Placenta % DNAm at 1st Trimester
Genomic Position on chr7
27169600
27170000
27170400
27170800
2717120
D
Endothelial
Hofbauer
Stromal
Trophoblasts
Villi
0/19
1/19
0/19
3/19
NA
Placenta Cell Type % DNAm at 1st Trimester
Genomic Position on chr7

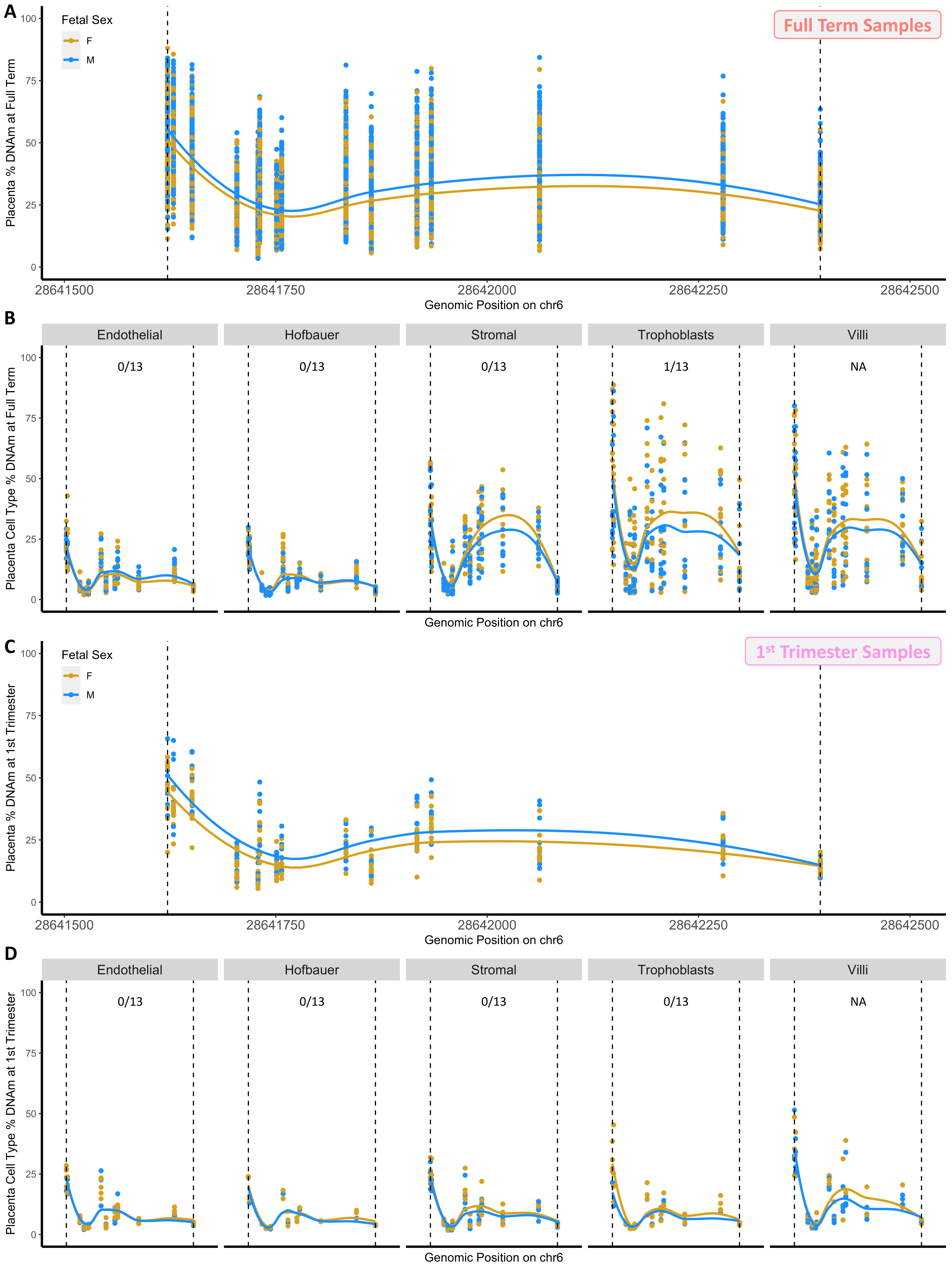

A
Fetal Sex
F
M
Full Term Samples
Placenta % DNAm at Full Term
100
75
50
25
0
28641500
28641750
28642000
28642250
28642500
Genomic Position on chr6
B
Endothelial
Hofbauer
Stromal
Trophoblasts
Villi
0/13
0/13
0/13
1/13
NA
Placenta Cell Type % DNAm at Full Term
Genomic Position on chr6
C
Fetal Sex
F
M
1st Trimester Samples
Placenta % DNAm at 1st Trimester
28641500
28641750
28642000
28642250
28642500
Genomic Position on chr6
D
Endothelial
Hofbauer
Stromal
Trophoblasts
Villi
0/13
0/13
0/13
0/13
NA
Placenta Cell Type % DNAm at 1st Trimester
Genomic Position on chr6
