## Supplementary material for "Large-scale placenta DNA methylation mega-analysis reveals fetal sex-specific differentially methylated CpG sites and regions": Description of Supplementary Data

**Supplementary Data 1: Full genome-wide association statistics for single site fetal sex differential methylation analysis in full term placenta samples.** Results from linear model regressing M-values onto estimated surrogate variables. Shown is output of topTable() function from the *limma* R package, with genomic position information of 450K probes appended.

**Supplementary Data 2: Full Gene Ontology enrichment results for genome-wide significant CpG sites from full term analysis.** Evaluated via gometh() function from the *missMethyl* R package.

**Supplementary Data 3: Full Gene Ontology enrichment results for genome-wide significant CpG sites from full term analysis, restricted to sites that do not distinguish between placenta cell types at term.** Evaluated via gometh() function from the *missMethyl* R package. Cell type discriminating probes determined by Yuan et al.

**Supplementary Data 4: Full Gene Ontology enrichment results for genome-wide significant CpG sites from full term analysis, restricted to sites that distinguish between placenta cell types at term.** Evaluated via gometh() function from the *missMethyl* R package. Cell type discriminating probes determined by Yuan et al.

**Supplementary Data 5: Full genome-wide association statistics for single site fetal sex differential methylation analysis in 3rd trimester placenta samples.** Results from linear model regressing M-values onto estimated surrogate variables. Shown is output of topTable() function from the *limma* R package, with genomic position information of 450K probes appended.

**Supplementary Data 6: Full genome-wide association statistics for single site fetal sex differential methylation analysis in 2nd trimester placenta samples.** Results from linear model regressing M-values onto estimated surrogate variables. Shown is output of topTable() function from the *limma* R package, with genomic position information of 450K probes appended.

**Supplementary Data 7: Full genome-wide association statistics for single site fetal sex differential methylation analysis in 1st trimester placenta samples.** Results from linear model regressing M-values onto estimated surrogate variables. Shown is output of topTable() function from the *limma* R package, with genomic position information of 450K probes appended.

**Supplementary Data 8: Full genome-wide association statistics for region-based fetal sex differential methylation analysis in full term placenta samples.** Results from *bumphunter* method with M-values and adjusted for estimated surrogate variables, implementing bootstrap-based multiple testing correction.

**Supplementary Data 9: Full genome-wide association statistics for region-based fetal sex differential methylation analysis in 3rd trimester placenta samples.** Results from *bumphunter* method with M-values and adjusted for estimated surrogate variables, implementing bootstrap-based multiple testing correction.

**Supplementary Data 10: Full genome-wide association statistics for region-based fetal sex differential methylation analysis in 2nd trimester placenta samples.** Results from *bumphunter* method with M-values and adjusted for estimated surrogate variables, implementing bootstrap-based multiple testing correction.

**Supplementary Data 11: Full genome-wide association statistics for region-based fetal sex differential methylation analysis in 1st trimester placenta samples.** Results from *bumphunter* method with M-values and adjusted for estimated surrogate variables, implementing bootstrap-based multiple testing correction.
